# Mechanistic modeling suggests that low-intensity focused ultrasound can selectively recruit myelinated or unmyelinated nerve fibers

**DOI:** 10.1101/2020.11.19.390070

**Authors:** Théo Lemaire, Elena Vicari, Esra Neufeld, Niels Kuster, Silvestro Micera

## Abstract

Low-Intensity Focused Ultrasound Stimulation (LIFUS) holds promise for the remote modulation of neuronal activity, but an incomplete mechanistic characterization hinders its clinical maturation. Here, we developed a computational framework to model intramembrane cavitation in multi-compartmental, morphologically-realistic neuronal representations, and used it to investigate ultrasound neuromodulation of peripheral nerves by spatially-varying pressure fields. Our findings show that LIFUS offers distinct parametric sub-spaces to selectively recruit myelinated or unmyelinated axons and modulate their spiking activity over physiologically relevant regimes and within safe exposure limits. This singular feature, explained by fiber-specific differences in membrane electromechanical coupling, consistently explains recent empirical findings and suggests that LIFUS can preferentially target nociceptive and sensory fibers to enable peripheral therapeutic applications not addressable by electric stimulation. These results open up new opportunities for the development of more selective and effective peripheral neuroprostheses. Our framework can be readily applied to other neural targets to establish application-specific LIFUS protocols.

## Introduction

Ultrasound (US)-based approaches have been increasingly adopted over the past decades for a variety of noninvasive therapeutic interventions (Escoffre and Bouakaz, 2016). These therapies rely on the mechanical nature of acoustic waves that propagate efficiently through biological tissue and can be accurately steered to concentrate mechanical energy within small volumes (∼mm3) around deep anatomical targets. In recent years, several *in vitro* and *in vivo* studies have shown that such acoustic waves can also be used to reversibly modulate the activity of various neural targets with remarkable spatial accuracy (Blackmore et al., 2019). These findings have propelled the development of low-intensity focused ultrasound stimulation (LIFUS) as a novel technology to achieve noninvasive, selective and reversible neuromodulation of virtually any neural structure.

Yet, despite a decade of intense investigation, several open issues have impeded the development of LIFUS as a clinically relevant technology. The variety of known physical effects of acoustic waves in biological tissue implies a wide range of possibilities for how neurons may translate mechanical energy into electrical responses, including membrane piezoelectricity (Heimburg and Jackson, 2005), flexoelectricity (Petrov, 2002) and mechanosensitive channels activation (Tyler, 2011). At the same time, distinguishing these candidate mechanisms in experimental settings and establishing their predominance over the multi-dimensional LIFUS parameter space remains a challenge. Consequently, it is difficult to provide a mechanistic perspective that would clarify and guide the heterogeneous and sometimes conflicting collection of neuromodulatory effects (excitatory and inhibitory, short and long term, localized and large-scale, reversible and permanent) obtained across animal models, neural targets, and experimental designs.

In light of these challenges, computational approaches have become helpful tools to increase the understanding of LIFUS-neuron interactions, as they allow a specific candidate mechanism to be examined. A significant effort made by Plaksin et al., who introduced the ***N**euronal **I**ntramembrane **C**avitation **E**xcitation* (NICE) model, described a candidate mechanism in which LIFUS induces the cavitation of specific phospholipidic structures (so-called bilayer sonophores), thereby dynamically altering membrane capacitance and triggering action potentials. This model predicts cell-type-specific LIFUS responses of cortical and thalamic neurons (Plaksin et al., 2016) that correlate indirectly with a range of empirical results obtained in the central nervous system (CNS) (Kim et al., 2012; King et al., 2013; Tufail et al., 2011; Yoo et al., 2011).

The NICE model, however, entails a significant numerical stiffness that has so far limited its applications to point-neuron studies (Plaksin et al., 2014, 2016; Tarnaud et al., 2018a) that could not address physiologically relevant questions, such as the influence of intracellular axial coupling and morphological inhomogeneity on neuronal responses, the spatiotemporal dynamics of those responses, and the impact of spatial features of the acoustic field on excitability (as is the case for electrical stimulation). Hence, a multi-compartmental model of intramembrane cavitation incorporating morphological details would be highly beneficial to increase our understanding of LIFUS neuromodulation by intramembrane cavitation in a more realistic setting.

In a recent study, we developed a *multi-**S**cale **O**ptimized **N**euronal **I**ntramembrane **C**avitation* (SONIC) model that alleviates the numerical stiffness of the NICE model by integrating the coarse-grained evolution of effective electrical variables as a function of a pre-computed, cycle-averaged impact of the oscillatory mechanical system (Lemaire et al., 2019), thereby drastically reducing computational costs while maintaining numerical accuracy. Building on this effective paradigm, we present *morphoSONIC*, a novel framework to simulate intramembrane cavitation into morphologically realistic neuron models. This framework leverages the optimized modeling and numerical integration pipelines of the NEURON simulation environment (Hines and Carnevale, 1997), and provides an alternative implementation of its internal cable representation as a hybrid (charge and voltage casted) electrical circuit that is numerically compatible with the SONIC model.

Specifically, we exploit this framework to investigate the mechanisms of ultrasonic neuromodulation in myelinated and unmyelinated peripheral fibers, using previously validated single-cable axon models (Reilly et al., 1985; Sundt et al., 2015). First, we provide an in-depth analysis of predicted LIFUS neuronal responses and recruitment mechanisms in both fiber types. Second, we characterize their LIFUS excitability by evaluating strength-duration (SD) “signatures” across a wide range of model and stimulation parameters and compare those signatures to those traditionally obtained with electrical stimulation. Third, we identify key morphological features underlying the distinct LIFUS sensitivities of myelinated and unmyelinated axons. Finally, we analyze cell-type-specific neuronal responses upon repeated acoustic exposure and identify pulsing protocols yielding a robust modulation of spiking activity.

## Results

We investigated the mechanisms of US neuromodulation by intramembrane cavitation in myelinated and unmyelinated fibers using morphologically realistic single-cable axon models of each fiber type (Reilly et al., 1985; Sundt et al., 2015) (SENN and Sundt model, respectively, **Figure 1**). These neural structures represent privileged, accessible neuromodulation targets, for which multiple studies have demonstrated the paramount influence of cellular morphology on the resulting excitability by electrical fields (McNeal, 1976; Rattay, 1986). As such, they are natural candidates for the study of LIFUS effects in morphologically realistic models.

**Figure 1.**
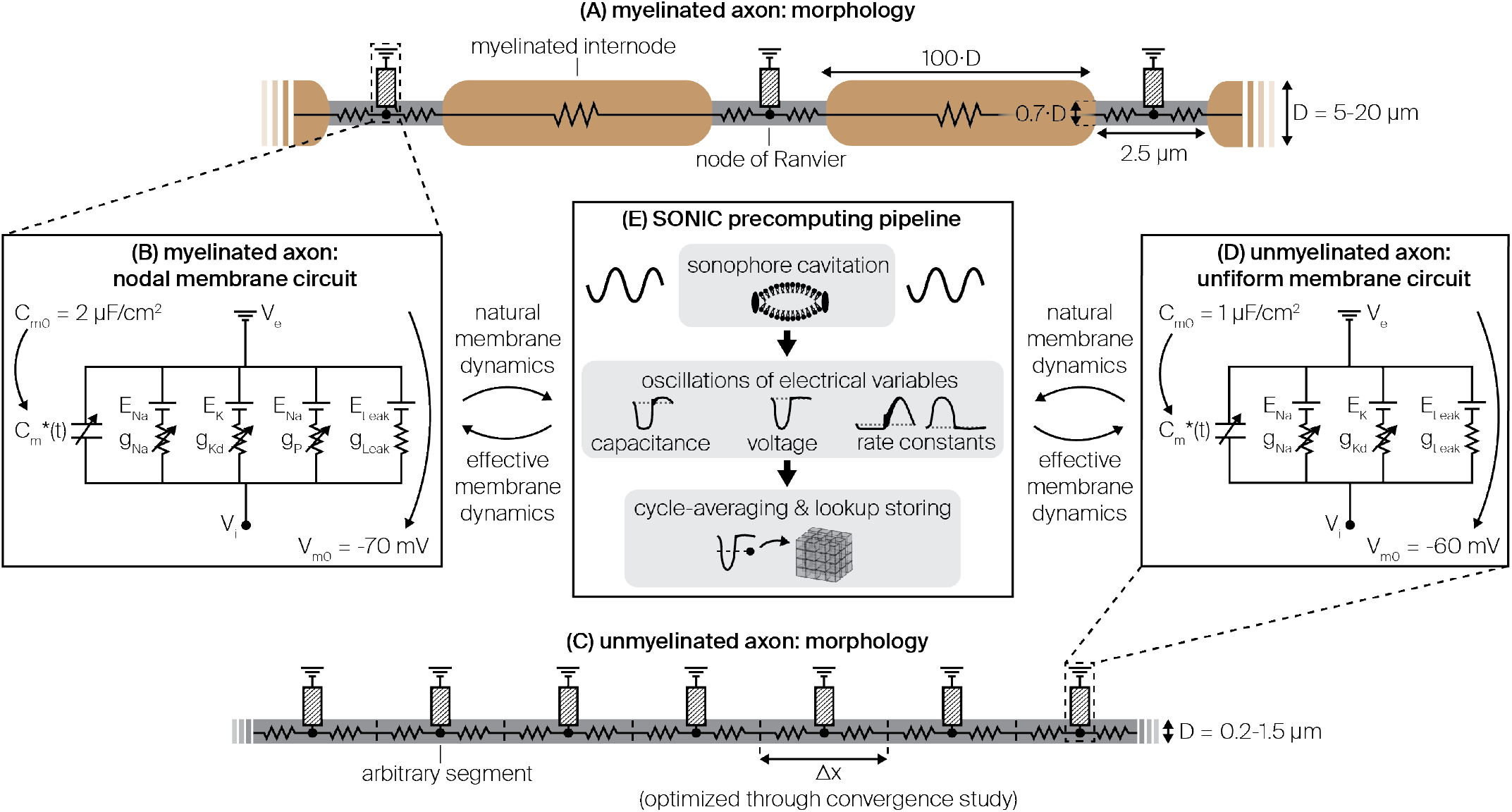
Morphology, biophysics and incorporation of the SONIC paradigm in myelinated and unmyelinated axon models. (A) Schematic of the myelinated axon model morphology. (B) Electrical circuit representation of the membrane dynamics at the nodes of Ranvier. (C-D) Equivalent morphological and biophysical descriptions of the uniform unmyelinated axon. (E) Schematic diagram

### The SONIC paradigm enables accurate simulations in multi-compartmental axon models

A recent study has shown that in multi-compartmental structures, the presence of large axial currents could introduce significant intra-cycle charge redistribution mechanisms, thereby inducing a significant divergence of the SONIC paradigm (Tarnaud et al., 2020). Thus, we aimed to establish the conditions of this divergence and whether it applies to the models of this study. To this end, we designed two-compartment benchmark models of intramembrane cavitation in which mechanical perturbation is modeled as a pure sinusoidal oscillation of membrane capacitance around its resting value with a specific amplitude in each compartment 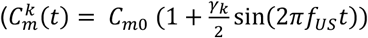, with *k* being the compartment index and γ_*k*_ the relative oscillation range, **Figure 2A**). This simplified perturbation produces neuronal responses in cortical models that are qualitatively comparable to those obtained with a detailed intramembrane cavitation model ((Plaksin et al., 2016), Fig. 9) and therefore facilitates the assessment of SONIC accuracy in the presence of a high-frequency, spatially-varying acoustic perturbation – arising either from pressure amplitude gradients or variations in sonophore coverage – without requiring the tedious integration of the mechanical model. Moreover, the sinusoidal drive can be intuitively mapped to original sonication parameters: for both axon models, relative capacitance oscillation ranges increased monotonically with acoustic pressure and showed little modulation by US frequency over the 20 kHz – 4 MHz range (**Figure 2B**), with rheobase excitation thresholds occurring around γ = 0.3 for the central, most-exposed node (see **Figure 5**).

**Figure 2.**
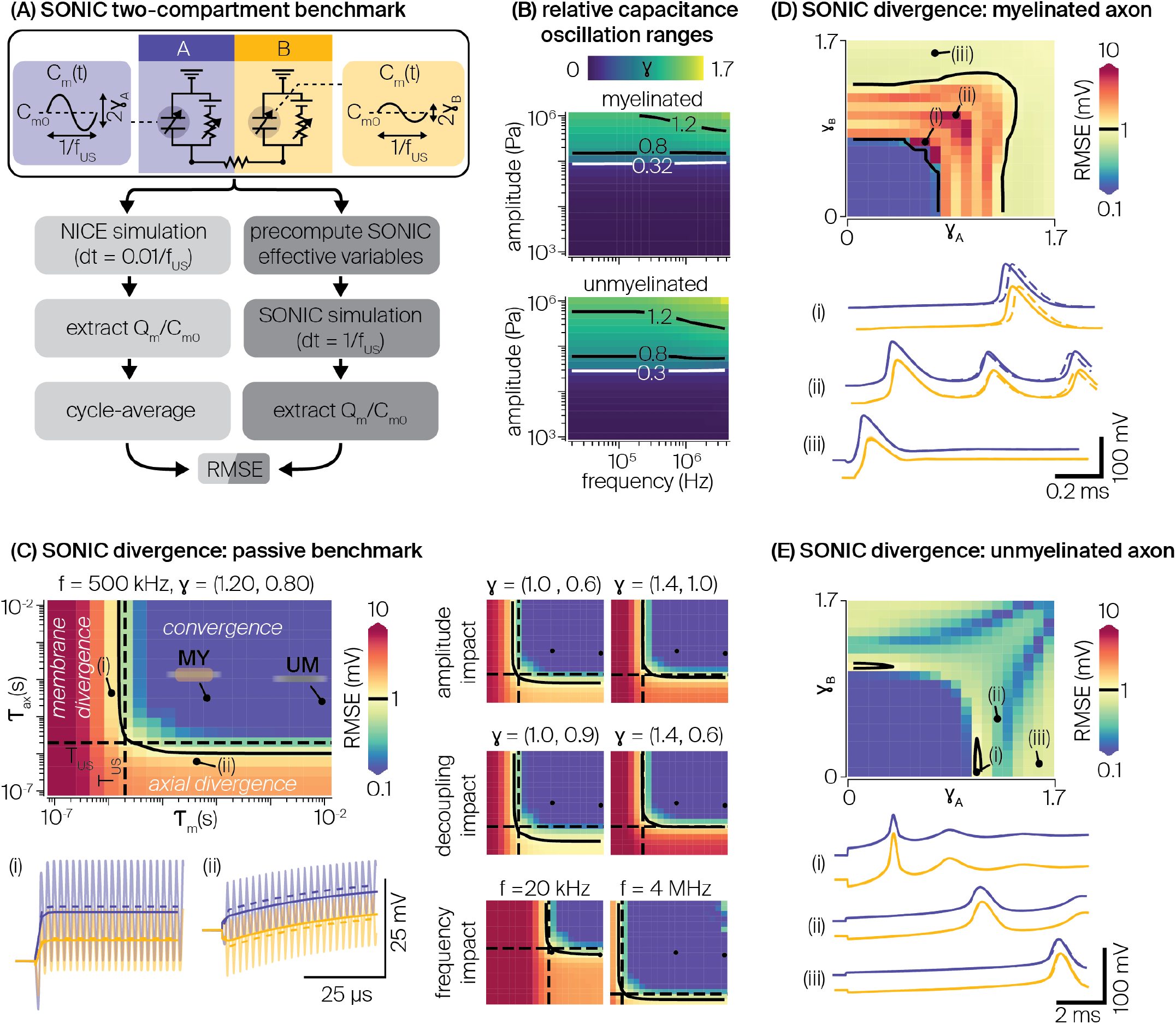
Evaluation of the SONIC paradigm accuracy in two-compartment benchmark models. (A) Schematic description of the two-compartment SONIC benchmark and associated divergence evaluation process. (B) Magnitude of the intra-cycle relative capacitance oscillation range as a function of stimulus frequency and pressure amplitude in myelinated and unmyelinated fibers, computed using axon-specific bilayer sonophore models at their respective resting charge density. White lines indicate cell-type-specific excitation threshold levels. (B) SONIC divergence (maximum RMSE between normalized charge density profiles resulting from SONIC and cycle-averaged NICE simulations) in a symmetric two-compartment passive model (C_m0_ = 1 μF/cm2, E_Leak_ = −70 mV), as a function of the model’s electrical time constants (τ_*m*_ = *C_m0_*/*g_Leak_*, *τ_ax_* = *C_m0_A_m_*/*G_a_*), for a typical US frequency (500 kHz) and sinusoidal oscillation drive (γ = (1.2, 0.8)). Acoustic periodicity, critical divergence level (1 mV), and fiber passive properties are indicated on the two-dimensional logarithmic color maps. Insets (i) and (ii) provide examples of NICE, cycle-averaged NICE, and SONIC charge density profiles for diverging conditions. Minimized maps evaluate SONIC divergence in the same model but for varying capacitance oscillation amplitudes and gradients, as well as varying US frequencies. (C) SONIC divergence in a two-compartment benchmark model of the myelinated axon for various combinations of oscillation pairs (f = 500 kHz, 1 ms stimulus). A critical divergence level (1 mV) is indicated on the color map. Insets (i-iii) provide examples of NICE, cycle-averaged NICE, and SONIC charge density profiles for various characteristic drive combinations. (D) Equivalent divergence evaluation in a two-compartment benchmark model of the unmyelinated axon (10 ms stimulus).

First, we investigated the influence of model and stimulus properties on the accuracy of the SONIC paradigm in predicting sub-threshold depolarization – a critical aspect of neuronal responses – using a passive benchmark model (see Methods). We evaluated SONIC divergence for various combinations of model membrane and axial time constants, providing quantitative estimates of the time taken by leakage and axial currents to respond to variations in transmembrane and longitudinal intracellular voltage gradients, respectively.

For a typical driving frequency (500 kHz) and supra-threshold oscillation ranges (γ = [1.2, 0.8]), the divergence of the SONIC paradigm showed a rather symmetrical dependency on both axial and membrane conductances (**Figure 2C**). Strong electrical conductances (i.e., short time constants) increase the sensitivity of the electrical system to the oscillatory mechanical drive to a point that currents instantaneously “convert” part of the capacitive displacement energy into fast-charge redistribution during an acoustic period, thereby impacting the net charge variation over that period. We can differentiate two distinct mechanisms of intra-cycle charge redistribution. On the one hand, leakage currents opposing intra-cycle deviations from reversal potentials create a transmembrane charge redistribution that reduces the net charge increase at each cycle, hindering the slow scale charge build-up in each compartment (**Figure 2C**, inset (i)). On the other hand, intracellular currents opposing intra-cycle voltage gradients create an axial charge redistribution across the two compartments that reduces the net charge gradient achieved over a cycle, ultimately limiting the magnitude of effective charge density gradients over time (**Figure 2C**, inset (ii)). Neither of these redistribution mechanisms were captured by the SONIC paradigm, which resulted in a divergent sub-space (*∈_max_* > 1 mV) for strong enough conductances. Conversely, weaker conductances (i.e. larger time constants) ensured minimal intra-cycle charge redistribution and outlined a sub-space of SONIC convergence (*∈_max_* < 1 mV). A clear bifurcation between these two sub-spaces emerged as the membrane and / or axial time constant approached the order of magnitude of the acoustic period.

Interestingly, the accuracy of the SONIC paradigm also depended on intrinsic stimulus features. On the one hand, both stronger amplitudes and larger gradients of capacitance oscillations amplified intra-cycle charge redistributions (and SONIC divergence) within the divergence sub-space, but they did not significantly affect the bifurcation time constant. Whereas, increasing oscillation frequencies expanded the convergence sub-space to shorter time constants, but did not significantly affect the magnitude of SONIC divergence within the divergence sub-space. Together, these findings suggested that the critical condition for SONIC convergence is that the model’s time constants should be longer than the drive periodicity ({τ_*m*_, τ_*ax*_} > 1/*f_US_*). The passive properties of both axon models used in this study satisfied this criterion, except at very low drive frequencies (f = 20 kHz).

Second, we investigated the applicability of the SONIC paradigm for the particular axon models used in this study. For this, we used two-compartment models with axon-specific morphological properties and full membrane dynamics, and evaluated SONIC divergence across a symmetric two-dimensional space of capacitance oscillation pairs with a drive frequency of 500 kHz.

As expected, limited drive oscillation amplitudes triggered passive build-ups in charge density in both models that were accurately captured by the SONIC paradigm (**Figure 2D-E**). As the capacitance oscillation amplitude reached a critical threshold in one compartment (*γ_thr_* = 0.7 and 0.9 for myelinated and unmyelinated benchmarks, respectively), both models then transitioned towards an active response. A few regions of SONIC inaccuracy appeared in the myelinated case around the transition threshold, where the timing of the action potential (AP) generation was especially sensitive to the sub-threshold build-up dynamics and therefore tended to amplify subtle differences in initial build-ups between the two paradigms (**Figure 2D**, insets (i)-(ii)). However, those differences vanished at higher oscillation amplitudes which elicited a more robust spiking dynamics (faster build-up, reduced post-spike oscillations) (**Figure 2D**, inset (iii)). It is also worth noting that SONIC inaccuracies were restricted to very few combinations of capacitance oscillation amplitudes, which corresponded to a narrow range of stimulus amplitudes. In the unmyelinated case, the slower intrinsic membrane dynamics and limited axial coupling enabled a robust SONIC accuracy across the entire explored oscillation range (**Figure 2E**, insets (i)-(iii)).

Taken together, these findings suggested that the SONIC model can provide accurate predictions of neuronal responses in single cable peripheral axon models across the entire LIFUS parameter space with the exception of very low US frequencies (f < 100 kHz) and can thus be reliably applied to investigate intramembrane cavitation in those models.

### Exogenous acoustic and electric sources produce normal field distributions along fibers

The simulation of spatially extended morphological models by artificial exogenous fields traditionally requires the spatial distribution of a perturbation variable across the model’s compartments to be solved. Thus, we aimed to analyze the nature of acoustic pressure and extracellular voltage distributions obtained from simplified, yet realistic configurations.

First, we used the distributed point source method (DPSM, see Methods) to predict acoustic propagation from a single element planar transducer in a water-like medium (Yanagita et al., 2009). Normalized pressure distributions in the propagation plane showed a high degree of directionality, with significant amplitudes concentrating along the main lobe normal to the transducer surface (Figure S1A). Both larger transducer diameters and higher frequencies induced a more pronounced near-field effect and shifted the focal distance further away from the source. At their respective focal distance, all configurations produced Gaussian-like pressure distributions along the transverse axis (Figure S1B), as confirmed by D’Agostino and Pearson’s statistical normality test on all distributions. Moreover, the full width at half maximum (FWHM) of these distributions increased linearly with the transducer radius and showed little modulation by US frequency (Figure S1C).

Second, used a point-source electrode approximation (see Methods) to predict extracellular voltage fields in a nerve-like environment (Ranck and Bement, 1965). Resulting distributions also showed some degree of directionality due to the medium anisotropic properties (Figure S1D). Distributions were somewhat simpler than for the acoustics case, with voltage amplitude decreasing as a function of distance from the electrode as a result of the increasing resistive path. Nevertheless, voltage distributions along the transverse axis were also “Gaussian-like”, and their width linearly increased as a function of the distance from the electrode (Figure S1E).

These results obtained using realistic propagation models show that both ultrasonic and electrical sources produce normal distributions of the perturbation variable along the fiber’s longitudinal axis, and that the “width” of those distributions can be controlled with a single parameter (transducer radius and electrode distance, respectively). Therefore, to assess the impact of stimulus spatial distribution on fiber excitability in a controlled manner and allow a direct comparison across the two simulation modalities, we sampled exogenous fields directly from Gaussian distributions of varying widths and amplitudes for the study of single, isolated fibers. In the following, the FWHM of the field distribution is referred to as the stimulus “beam width”.

### LIFUS modulates membrane capacitance to excite myelinated and unmyelinated axons

In this section, we describe “typical” predicted responses of myelinated and unmyelinated axons to ultrasound stimulation. We selected standard axon models using representative diameters for each axon population (10 μm and 0.8 μm for myelinated and unmyelinated fibers, respectively), and assumed a uniform sonophore depiction across the model non-insulated compartments (i.e. Sundt unmyelinated membrane and SENN Ranvier nodes). We chose a typical sonophore radius (a = 32 nm) used in previous studies (Lemaire et al., 2019; Plaksin et al., 2016, 2014) and a physiologically plausible sonophore coverage fraction (f_s_ = 80%) falling within a range of conserved excitability in cortical point-neuron models (see (Lemaire et al., 2019), Fig. 10). Importantly, those parameters were chosen ad-hoc, without re-tuning or post-hoc adjustments. We also considered “standard” acoustic perturbations, using a typical US frequency (500 kHz) and sampling acoustic pressures stemming from a 5 mm-wide Gaussian distribution (i.e. qualitatively equivalent to the distribution generated by a 10 mm diameter single-element planar transducer at focal distance at this frequency) with a peak amplitude of 120 kPa (i.e. significantly above excitation thresholds of both axons). Finally, we examined responses to pulse durations for which the rheobase regime is approached for each fiber type (1 ms and 10 ms for the myelinated and unmyelinated fibers, respectively, see next section). To better identify mechanisms of axonal recruitment by US pulses, we quantified the time required to reach a normalized charge build-up of 5 mV in the central compartment for each response and computed the contribution of each individual current to this initial build-up.

For both models, the sonication pulse onset generated instantaneous drops in effective membrane capacitance in the axon compartments, whose magnitude increased with the amplitude of the local acoustic pressure, thereby amplifying the absolute value of transmembrane voltage and inducing hyperpolarization (**Figure 3A-B**). Due to the Gaussian distribution of acoustic pressure along the axon, central compartments experienced a stronger hyperpolarization than peripheral ones, which introduced a longitudinal gradient in transmembrane voltage along the fiber.

**Figure 3.**
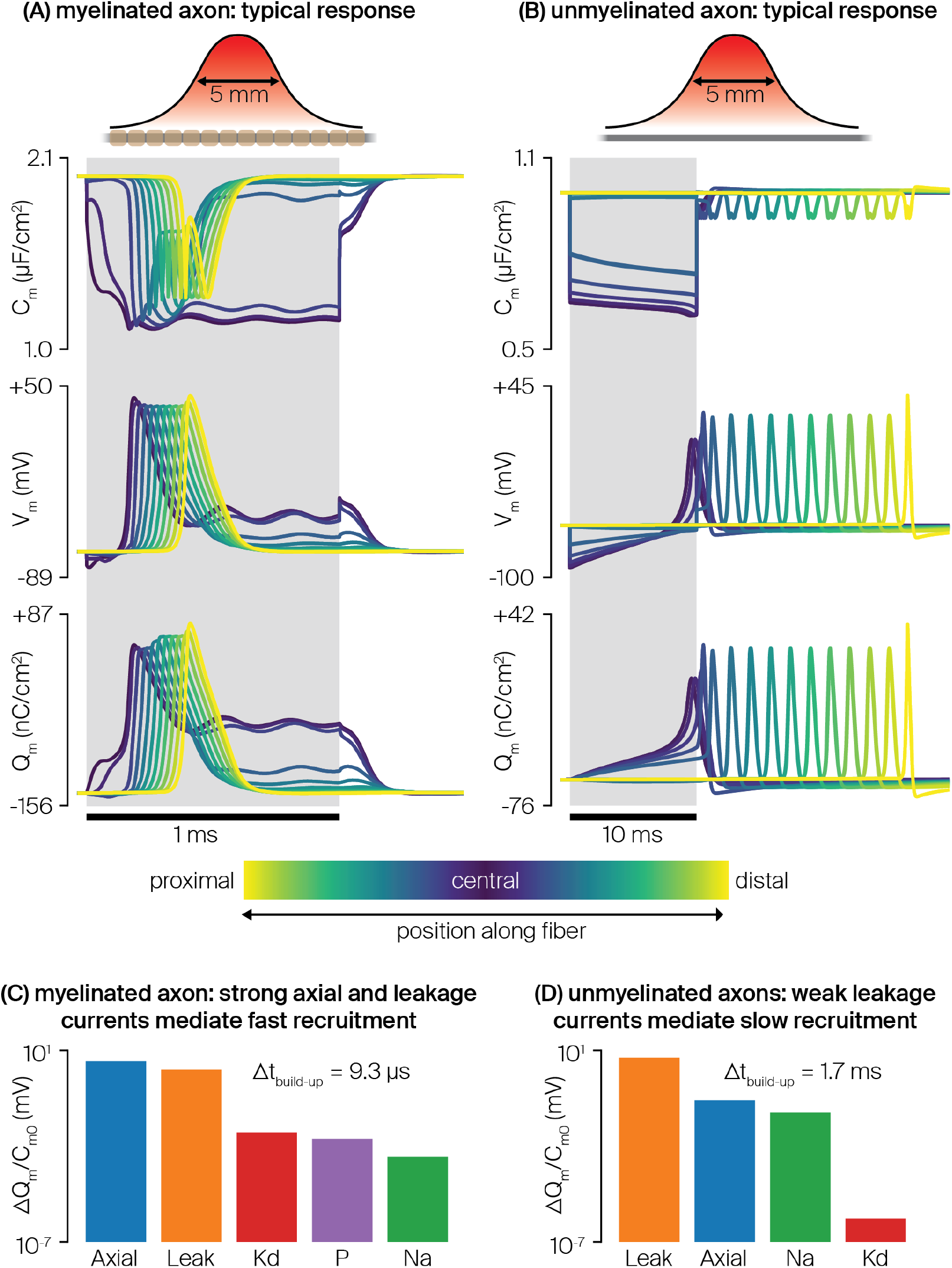
Typical responses of myelinated and unmyelinated axon models to a single US pulse. (A) Time profiles of effective membrane capacitance, effective membrane potential, and effective membrane charge density across compartments during a typical response of a myelinated axon to a 1 ms sonication (500 kHz frequency, 5 mm-wide Gaussian pressure distribution aligned on the fiber with a spatial peak of 120 kPa). (B) Equivalent time profiles during the typical response of an unmyelinated axon to a 10 ms sonication (identical pressure distribution as in (A)). (C) Quantification of the membrane and axial currents contributions to the first 5 mV of normalized charge build-up in the fiber central compartment. (D) Equivalent quantification for the response of the unmyelinated axon’s central compartment.

At the central compartment where hyperpolarization is the largest, leakage currents arose to bring transmembrane voltage towards the leakage reversal potential, thereby inducing a build-up in local charge density. The considerably higher density of leakage channels in the Ranvier node compared to the unmyelinated membrane 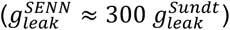 induced much larger leakage currents (**Figure 3C-D**). Moreover, significant axial currents also arose in the myelinated axon, driven by large voltage gradients between the central and neighboring Ranvier nodes. Together, these two depolarizing currents yielded a much faster membrane charge build-up in the myelinated axon, yielding shorter response latencies. These differences are reflected in the times required to achieve a normalized charge build-up of 5 mV in the central compartment (9 μs and 1.7 ms for myelinated and unmyelinated fibers, respectively).

For a long enough sonication, membrane charge density increased until a spiking threshold was reached, prompting the opening of Sodium ion channels and thereby triggering an AP in the central compartment that started travelling bi-directionally towards the axon extremities. As expected, both axons exhibited marked differences in conduction velocities: fast saltatory conduction in the large diameter myelinated axon allowed the AP to reach the extremities of the axon in less than 1 ms, whereas that process took more than 10 ms in the slowly conducting unmyelinated axon. As the sonication outlasted the AP duration in the myelinated axon, affected nodes transitioned into a “plateau potential” regime (stabilization of membrane charge density around a depolarized value).

Finally, the sonication offset removed the mechanical membrane perturbation, and effective membrane capacitances instantaneously reverted to their resting values, triggering a rapid reduction in transmembrane voltage magnitudes. The myelinated axon then simply repolarized back to its equipotential resting state, while the AP propagated towards peripheral extremities in the unmyelinated axon.

The effect of electro-mechanical coupling was visible across neuronal responses. During the sub-threshold charge build-up, the decrease in electrical pressure (a constraining force on the bilayer sonophore, proportional to 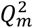) amplified membrane deflections, which further reduced the effective membrane capacitance (an effect more pronounced on the myelinated axon). In addition, the propagating spike induced a wave of time-varying electrical pressure that also modulated the effective membrane capacitance.

### LIFUS can selectively recruit myelinated and unmyelinated axons

In the previous section, we showed that ultrasonic axon recruitment requires the membrane charge density to be brought locally above a spiking threshold to engage voltage-gated channels. Yet, the underlying mechanisms eliciting this charge build-up differ significantly from those of electrical stimulation (McNeal, 1976; Rattay, 1986). Therefore, we aimed to determine if the two stimulation modalities could produce distinct excitability patterns. To this end, we computed excitation thresholds for various pulse durations ranging from 10 μs to 1 s (using binary search procedures) to construct strength-duration (SD) curves.

First, we evaluated the excitability of representative myelinated and unmyelinated axons (10 μm and 0.8 μm diameters, respectively) with a typical stimulus width (5 mm). With electrical stimulation, threshold peak extracellular voltages required to elicit a travelling AP decreased with increasing pulse duration, and then reached an asymptotical (so-called “rheobase”) regime for long enough pulses (**Figure 4A**). In line with previous modeling studies (Lubba et al., 2019; Tarnaud et al., 2018b), excitation thresholds for the myelinated axon were lower than those of the unmyelinated axon over the entire range of pulse durations. This result can be explained by two main factors. For short pulses where the speed of the depolarization predominantly determines when / if the spiking threshold is reached, myelinated axons can be recruited because of short membrane time constants, whereas unmyelinated axons fail to respond fast enough. Conversely, for long pulses approaching the rheobase regime, transient features become less critical and longitudinal gradients of the applied extracellular voltage become the main determinant of axonal excitability (Warman et al., 1992). Here again, myelinated axons are easier to recruit because of their insulated internodes that effectively discretize the voltage field at sparsely distributed Ranvier nodes, thereby producing stronger longitudinal gradients at the central node than those encountered across the continuous membrane of unmyelinated axons. With ultrasonic stimulation, threshold peak acoustic pressure amplitudes required for excitation also decreased with increasing pulse durations and reached a rheobase regime for long enough pulses (**Figure 4B**). Similarly as with electrical stimulation, short membrane and axial time constants conferred a low response latency to myelinated axons (see **Figure 3**), thereby allowing excitation by short ultrasonic pulses to which unmyelinated axons failed to respond. Surprisingly however, for longer pulse durations (*PD* ≥ 10 mm), the SONIC paradigm predicted lower excitation thresholds in unmyelinated axons than in myelinated axons.

**Figure 4.**
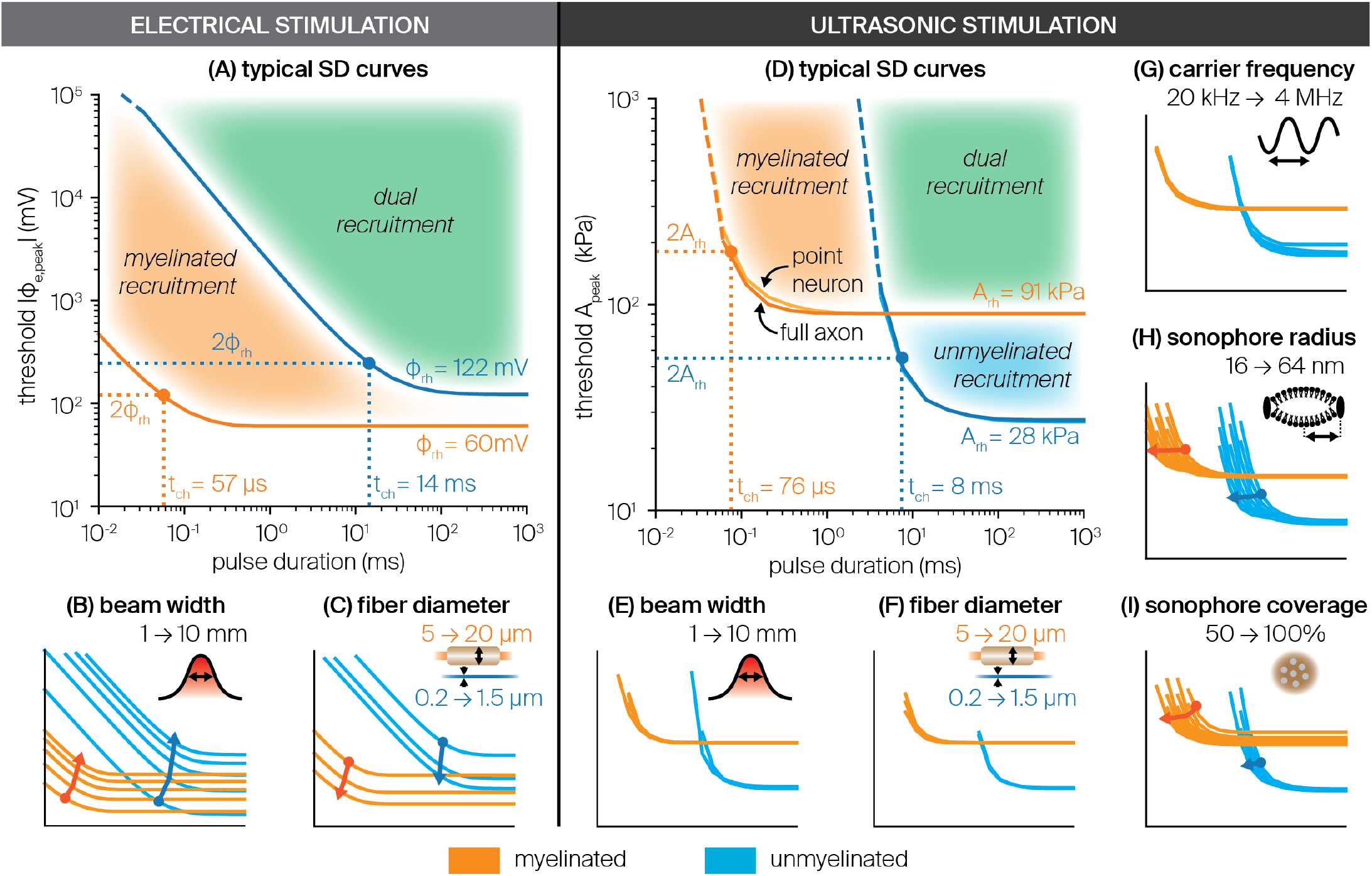
Comparison of strength-duration curves of myelinated and unmyelinated axons upon electrical and ultrasonic stimulation. (A) SD curves of representative myelinated (10 μm diameter, in orange) and unmyelinated (0.8 μm diameter, in blue) axons, depicting the threshold absolute peak extracellular voltage required to elicit fiber excitation as a function of pulse duration, for a characteristic 5 mm wide Gaussian extracellular voltage distribution. Rheobase and chronaxie values of each curve are indicated, as well as distinct areas of fiber recruitment. (B) SD curves of representative myelinated and unmyelinated axons for Gaussian extracellular voltage distributions of varying widths (1 to 10 mm). Arrows indicate the translation of the chronaxie point in the SD space for increasing stimulus width. (C) SD curves both fiber types of upon stimulation with a characteristic voltage distribution, for varying fiber diameters within the physiological range of each population (myelinated: 5 to 20 μm, unmyelinated: 0.2 to 1.5 μm). Arrows indicate the translation of the chronaxie point in the SD space for increasing fiber diameter. (D) SD curves of representative myelinated and unmyelinated axons, depicting the threshold peak acoustic pressure amplitude required to elicit fiber excitation as a function of pulse duration, for a characteristic 5 mm wide Gaussian acoustic pressure distribution and US frequency (f_US_ = 500 kHz), using typical values of sonophore radius (a = 32nm) and sonophore coverage fraction (f_s_ = 80 %) in the model’s compartments. SD curves using equivalent “node” models located under the stimulus peak are also indicated (light blue and orange curves), as well as rheobase and chronaxie values of each curve, and distinct areas of fiber recruitment. (E) SD curves of representative myelinated and unmyelinated axons with typical US frequency and sonophore parameters for Gaussian pressure distributions of varying widths (1 to 10 mm). (F) SD curves both fiber types with typical US frequency, pressure distribution and sonophore parameters, for varying fiber diameters within the physiological range of each population. (G) SD curves of equivalent “node” models of both fiber types with typical sonophore parameters and pressure distributions for varying US frequencies (20 kHz to 4 MHz). (H) SD curves of “node” models with typical pressure distribution, US frequency and sonophore coverage fraction for varying sonophore radii (16 to 64 nm). (I) SD curves of “node” models with typical pressure distribution, US frequency and sonophore radius, for varying sonophore coverage fractions (50 to 100 %).

Second, we characterized the impact of stimulus beam width and fiber diameter on excitability by systematically exploring relevant parameter ranges and using the “chronaxie point” (i.e. the pulse duration at which the threshold is twice the rheobase, see **Figure 4A-B**) as a reference point to measure the rigid translation of SD curves in the (pulse duration – stimulus amplitude) space. With electrical stimulation, narrowing stimulus beam widths enhanced excitability of myelinated and unmyelinated axons by producing stronger longitudinal gradients in extracellular voltage (**Figure 4C**). These stronger gradients primarily translated SD curves towards lower thresholds, but they also slightly diminished chronaxie durations. Very narrow beams produced an inversion of rheobase values, and unmyelinated axons became easier to recruit with long enough pulses. In a mirroring manner, increasing fiber diameters also enhanced excitability in both axon types (**Figure 4D**) as a result of (i) a larger intracellular conductance amplifying depolarization in response to a given extracellular voltage gradient, and (ii) in myelinated axons, an increased internodal spacing (*L* = 100*D_fiber_*) that further amplifies longitudinal gradients between consecutive Ranvier nodes. Again, both of these effects induced considerable shifts of SD curves towards lower thresholds and slightly reduced chronaxie durations in both axon types. Conversely, with ultrasonic stimulation, SD curves were remarkably consistent across a range of stimulus beam widths, as well as across the physiological range of fiber diameters of both populations, with only very slight variations in the chronaxie point and no clear trend emerging (**Figure 4E-F**).

The relatively low sensitivity of ultrasonic excitation thresholds to stimulus beam width and fiber diameter suggest that excitability patterns are primarily dictated by the magnitude of the peak acoustic pressure along the axon, rather than by the beam shape or the axial properties of the axon. To verify that hypothesis, we carried out the same excitability analysis in point-neuron models representing isolated neuronal compartments of the two axon models, namely a SENN Ranvier node and a Sundt unmyelinated segment, referred to as “node” models. We found almost identical SD curves between the node and full axon models (**Figure 4D**), thereby confirming that excitation is primarily mediated by the localized action of acoustic pressure on the cellular membrane. At first glance, these results seem to challenge the observation that axial currents contribute significantly to the initial charge build-up at the central node of myelinated axons upon sonication (**Figure 3**), and may therefore indicate the presence of a sharp transition in the mechanical response of the membrane to intensifying acoustic fields, bringing axons from passive to active responses within narrow amplitude ranges. Nevertheless, these results suggest that LIFUS-triggered excitation is primarily a local phenomenon - at least in these models – that can be accurately predicted without considering extended morphological details.

Given the high accuracy of node models in predicting cell-type-specific excitation thresholds, we leveraged their computational efficiency to explore the impact of acoustic frequency, sonophore size and sonophore coverage on neuronal excitability. In line with previous modeling results in CNS neurons (Lemaire et al., 2019; Plaksin et al., 2014) we found that US frequency does not significantly affect excitation thresholds apart from a slight increase above 1 MHz due to higher viscous stresses limiting sonophore cavitation (**Figure 4G**). Moreover, increasing sonophore radii induced mainly a “horizontal” shift of excitability towards shorter durations (**Figure 4H**), while increasing sonophore coverage fractions reduced both threshold baselines and chronaxie durations (**Figure 4I**).

### Resting membrane capacitance governs fiber excitability for long pulse durations

Strength-duration analyses revealed that unmyelinated axons exhibited lower excitation thresholds for long ultrasonic pulses, a trend robust to variations in model and stimulus parameters. Thus, we aimed to investigate the underlying mechanisms supporting this enhanced excitability using cell-type-specific node models, which proved to be appropriate benchmark tools to study ultrasonic neuronal recruitment (**Figure 4**).

For sub-threshold acoustic amplitudes and rheobase pulse durations, both the myelinated and the unmyelinated nodes responded to sonication with a build-up in charge density towards a more depolarized steady state (**Figure 5A**). Increasing the acoustic amplitude enhanced the magnitude of this build-up until the node’s spiking threshold was reached and an AP was fired. Interestingly, the exponential convergence of sub-threshold charge build-ups indicated that they were mostly mediated by passive currents, and could therefore be approximated by a simple RC membrane circuit with a single leakage conductance. Under this approximation, the steady-state charge build-up is proportional to the variation of effective membrane capacitance 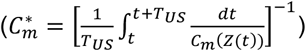 from its resting value:

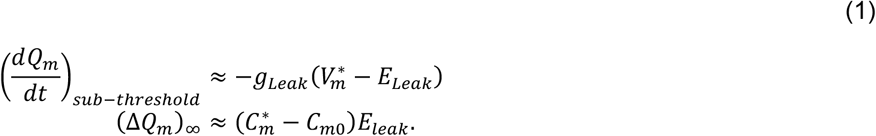

For both cell types, the passive circuit approximation can accurately predict the magnitude of steady-state charge build-up across a cell-type-specific range of sub-threshold acoustic amplitudes. This high prediction accuracy confirms that sub-threshold dynamics is almost entirely governed by the drop in effective capacitance.

**Figure 5.**
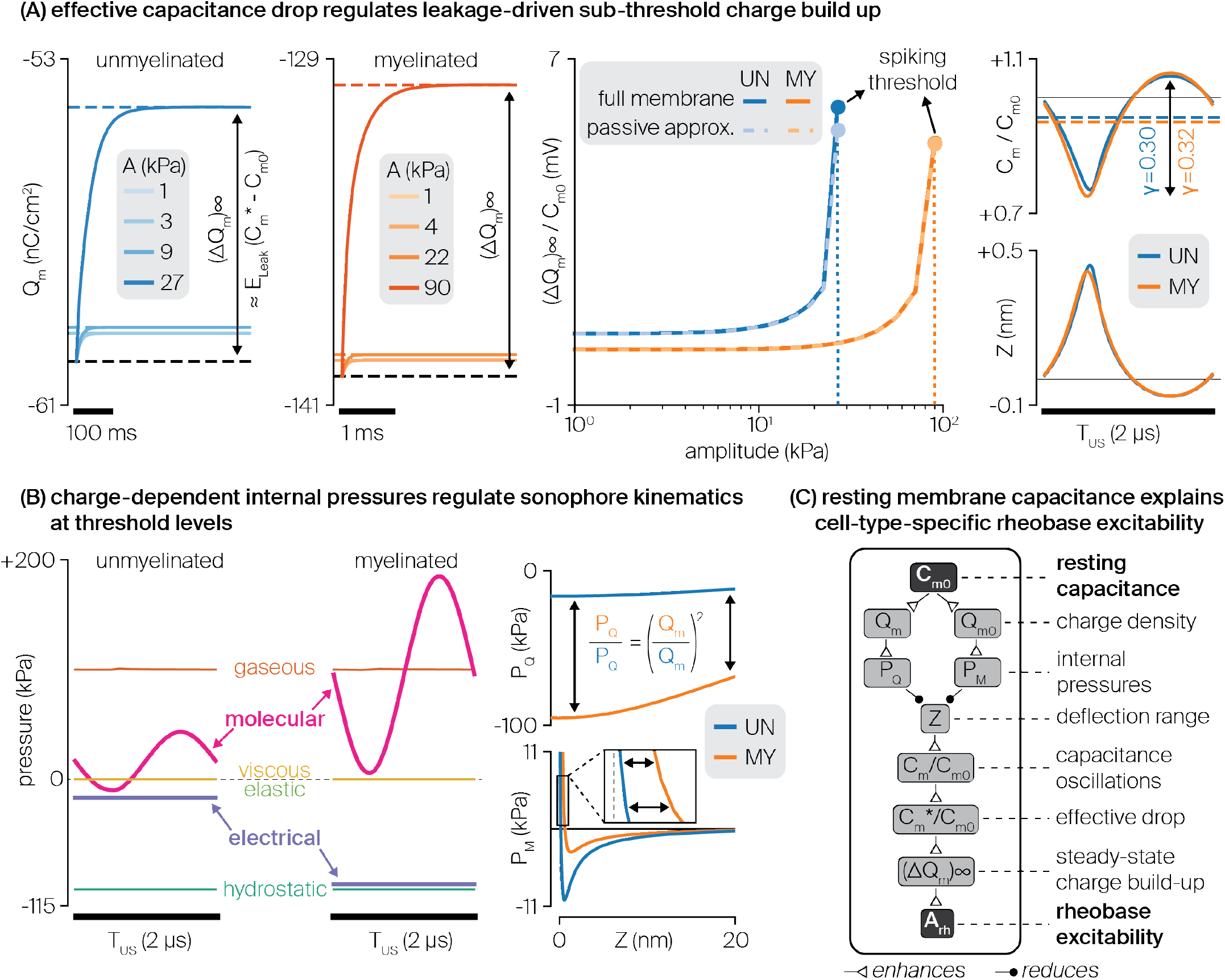
Underlying mechanisms of distinct rheobase excitabilities in myelinated and unmyelinated axons. (A) Effective capacitance variations regulate sub-threshold charge build-ups. From left to right: LIFUS-triggered, exponentially converging charge build-ups in myelinated and unmyelinated “node” models for various sub-threshold pressure amplitudes. Normalized steady-state charge build-ups for each “node” model as a function of sub-threshold pressure amplitude, computed from full membrane simulations (plain lines) and estimated from the sole relative variation in effective membrane capacitance (dashed lines, passive circuit approximation). Detailed intra-cycle oscillation profiles of membrane capacitance and membrane deflection for each fiber type at their respective threshold levels. (B) Charge-dependent electrical and molecular pressure regulate threshold sonophore kinematics. From left to right: detailed profiles of internal pressure forces regulating sonophore cavitation during an acoustic period, driven by cell-type-specific threshold acoustic pressures. Detailed profiles of electrical and molecular pressures in both fiber types along the physiological range of membrane deflection. (C) Schematic diagram showing the causal chain of influence by which resting membrane capacitance affects charge-dependent internal pressures, sonophore kinematics, effective capacitance variations, and ultimately rheobase excitability.

Both the myelinated and unmyelinated nodes required similar normalized charge build-ups to reach the spiking threshold (5.0 and 5.9 mV respectively), which corresponded to comparable relative variations in effective membrane capacitance (−6.3 and −5.2 %). Moreover, looking at intra-cycle dynamics, the resemblance of these effective cycle-averaged values arose from analogous oscillation profiles of membrane capacitance over an acoustic period (normalized oscillation ranges of 0.32 and 0.30, respectively). Recalling that capacitance is defined here as a deflection-dependent variable (see Materials and Methods section), this cross-model analogy could be mapped further back to cavitation profiles. Surprisingly, however, these similar membrane deflections were achieved at significantly different acoustic pressure amplitudes (91 kPa and 28 kPa, respectively). This discrepancy indicates variations in the internal kinetic system regulating sonophore cavitation dynamics in each node model.

Closer inspection of the detailed oscillation profile and resulting signal energy of each internal pressure component at these cell-type-specific amplitudes revealed several interesting features (**Figure 5B**). First, the relatively small cavitation magnitudes and velocities for these threshold levels 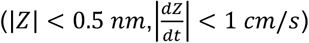 did not generate significant viscoelastic stresses on the membrane and surrounding medium. Moreover, this cavitation dynamics allowed for an instantaneous equilibration of gaseous and hydrostatic pressures on both sides of the sonophore cavity through transmembrane gas transport, thereby yielding identical energy levels for these pressure components across the two models. In contrast, both electrical and molecular pressures showed much larger energy levels for the myelinated sonophore model than for its unmyelinated counterpart. More specifically, the molecular pressure profile was shifted towards more positive values and showed higher oscillation amplitudes, whereas the electrical pressure profile was constant across a cycle but shifted towards more negative values. Together, these two pressure components are responsible for the cell-type-specificity of sonophore cavitation kinetics.

These changes in dynamic pressure oscillations can be mapped back to distinct profiles over a reference range of membrane deflections, allowing for the elucidation of the mechanisms of cell-type-specific rheobase excitability:

- The electrical pressure accounts for the attraction forces between the electric ion charges on the membrane leaflets, and is defined as 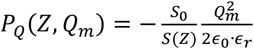. Therefore, both electrical pressure profiles show a weak dependence on membrane deflection, and a constant magnitude ratio across the deflection range 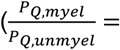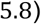, corresponding exactly to the square of the ratio of threshold charge densities across the two models. 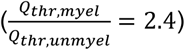The latter ratio primarily arises from variations in a fundamental biophysical property: the resting specific membrane capacitance of the myelinated axon is twice as high as that of the unmyelinated axon (see Methods). This increased capacitance allows the myelinated membrane to accumulate twice as much charges for identical transmembrane voltages, thereby increasing the electrical pressure on the membrane and hindering sonophore expansion around threshold levels.
- The intermolecular pressure is defined by a Lennard-Jones expression integrated across the sonophore surface: 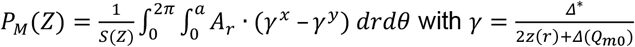. All parameters of this expression are fixed except for *Δ*^*^, the gap between the two membrane leaflets in the absence of charges. This parameter is calculated from a model-specific equilibrium state that depends on resting charge density, and therefore shows cell-type-specificity: the more negative resting charge density of the myelinated axon – mainly resulting from its larger capacitance – results in a smaller computed gap compared to the unmyelinated axon (1.1 nm vs 1.3 nm, respectively). Slight changes in this key parameter have profound implications on the resulting molecular pressure profiles: the smaller gap in the myelinated model reduces the amplitude of the negative (i.e. attractive) peak, and more importantly, shifts the transition towards positive (i.e. repulsive) pressure to a more positive deflection value, thereby producing much larger values of repulsive intermolecular pressure and hindering sonophore compression during an acoustic cycle around threshold levels.

The resting membrane capacitance is thus a crucial parameter that indirectly regulates the rheobase excitability of peripheral axons. This regulation is explained by a causal chain of influence (**Figure 5C**), can be summarized as follows: the resting capacitance influences both the resting value and the variation range of membrane charge density, thereby influencing charge-dependent internal pressures. That is, with larger capacitance, electrical pressure becomes more constraining during expansion phases and intermolecular pressure becomes more repulsive during compression phases. Together, these two pressure amplifications restrict the cavitation dynamics, and thus require higher acoustic pressures to attain similar membrane deflection and resulting relative capacitance oscillation ranges. In terms of cycle-averaged dynamics, higher pressures are needed to reach a given relative drop in effective capacitance, which almost entirely regulates the sub-threshold charge build-up. Given that both axon models require similar relative charge build-ups to reach their spiking threshold, rheobase excitability is then predominantly determined by the electrical modulation of cavitation dynamics, and hence by the resting membrane capacitance. In light of this mechanism, the enhanced excitability of unmyelinated axons for long pulse durations is explained by their smaller resting capacitance.

### Pulsed LIFUS robustly modulates axon spiking activity

In the previous sections, we analyzed response and excitability patterns of axon models upon application of isolated ultrasonic pulses. In the following, we investigated how the repeated application of such pulses can be used to modulate the spiking activity of axons over time. To do so, we simulated full axon models (using the standard model parameters defined in previous sections) upon the application of 10 consecutive sonication pulses (setting the stimulus beam width to one fifth of the fiber length), detected propagated APs on membrane charge density traces of axon extremities, and computed the resulting firing rate as the reciprocal of the average inter-spike interval over the simulation window.

We first evaluated the impact of pulsing parameters on spiking activity for a fixed acoustic pressure distribution with a peak amplitude of 300 kPa (a value falling safely above single pulse excitation thresholds of both axon models). Given the important differences in the LIFUS response time constants observed between myelinated and unmyelinated axons, we explored a relevant range of pulse durations around the axon’s single pulse chronaxie for each model.

Myelinated axons responded with very low latency but only fired a single spike for each acoustic pulse, followed by a stabilization to a plateau potential regime. As a result, they could be driven very robustly to follow the pulse repetition frequency (PRF) up to approximately 1 kHz over a wide range of pulse durations (**Figure 6A**, inset (i)). At higher stimulus rates, repeated pulses started to interfere with the cell’s refractory period, thereby preventing spike generation and / or propagation on average every two pulses, causing the axon to synchronize with the half-PRF (inset (ii)). At even higher stimulus rates, only very short pulses enabled a sustained firing activity, as the axon progressively reached its physiological limit at around 1 kHz (inset (iii)).

**Figure 6.**
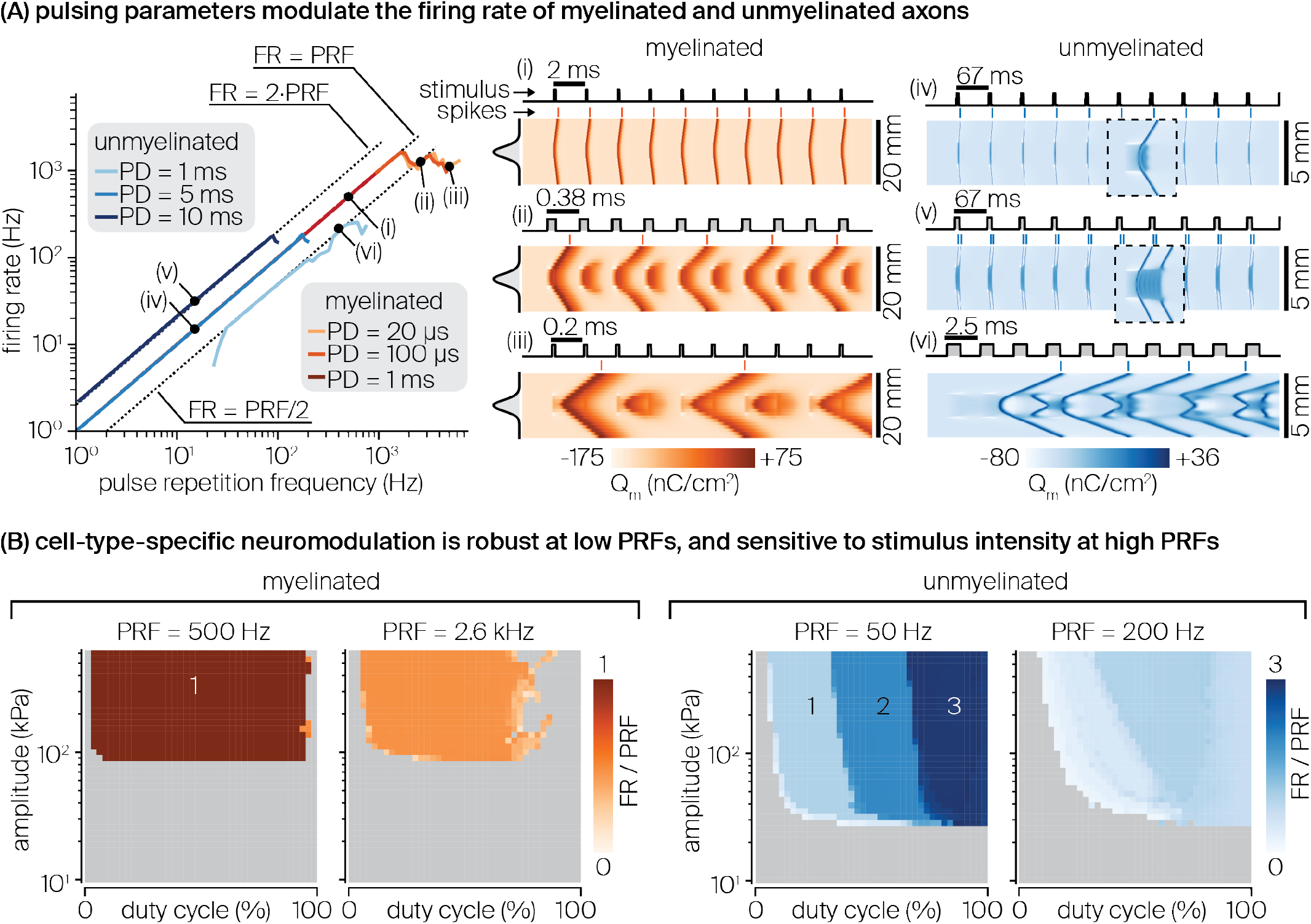
Modulation of spiking activity by pulsed sonication in myelinated and unmyelinated axons. (A) Average firing rate elicited in each axon type by a Gaussian acoustic pressure distribution covering one fifth of the fiber length, using default sonophore parameters (a = 32nm, f_s_ = 80%) and US frequency (fUS = 500 kHz), for various pulse durations and pulse repetition frequencies. Dashed lines indicate half, one time and double of the stimulus rate. Detailed spatiotemporal profiles of membrane charge density are indicated for characteristic spiking regimes of each fiber type, along with detailed profiles of the stimulus spatial distribution (vertical) and temporal application (horizontal). (B) Average firing rate (normalized by pulse repetition frequency) elicited in each fiber type as a function of duty cycle and peak acoustic pressure amplitude for cell-type-specific pulse repetition frequencies yielding “robust” and “sensitive” spiking behaviors. Numbers on the color maps indicate characteristic regimes of normalized firing rate.

In contrast, unmyelinated axons responded with higher latency, but could fire multiple spikes for prolonged sonication. Consequently, their behavior at low PRFs strongly depended on the pulse duration: short pulses (PD < 5 ms) did not induce any response, intermediate pulses (5 < PD < 10 ms) induced PRF-locking (inset (iv)), and long enough pulses (PD >= 10 ms) induced spiking activity at double or even higher multiples of the stimulus rate (inset (v)). At intermediate stimulus rates (20 Hz < PRF < 100 Hz), temporal summation of sub-threshold responses enabled recruitment by short pulses at half the PRF or below. Above 100 Hz, the range of available pulse durations was progressively restricted to shorter values that only allowed the fiber to fire at half the stimulus rate or below (inset (vi)), until a physiological limit was reached around a firing rate of 200 Hz.

Having established that pulsing parameters trigger cell-type-specific patterns of spiking activity, we aimed to investigate whether these neuromodulatory effects also depend on stimulus intensity. Hence, we simulated each axon type across a two-dimensional space of duty cycles (DC, from 0 to 100%) and peak pressure amplitudes (from 10 to 600 kPa), and for each combination, computed the resulting firing rates normalized by the PRF.

At low PRFs allowing a robust pulse-spike synchronization (identified from **Figure 6A** for each axon type), neuromodulatory effects were surprisingly consistent across a wide range of supra-threshold stimulus amplitudes (**Figure 6B**). The myelinated axon fired exactly one spike per pulse for *DC* ∈ [0.02, 0.95] (i.e. for pulses long enough to allow a first response yet distant enough to avoid destructive interaction with the refractory period), independently of stimulus amplitude. In contrast, the unmyelinated axon initiated a first response at slightly larger DCs and then exhibited three distinct spiking regimes with 1, 2 and 3 spikes per pulses as DC increased up to 1. A slight dependency on stimulus amplitude was noted here, as larger pressures shifted transitions between the different spiking regimes to lower duty cycles.

At high PRFs allowing only sub-stimulus rate spiking activity (see again **Figure 6A**), neuromodulatory effects were more intricate, and showed more dependency on stimulus amplitude. In this high-frequency regime (PRF = 2.6 kHz), the myelinated axon’s firing rate approached a maximum of 0.5 times the PRF over a wide duty cycle interval (*DC* ∈ [0.04, 0.70]). At larger duty cycles, spiking was only elicited for sparse DC-amplitude combinations allowing an optimal trade-off between fast depolarization to US stimuli and limited destructive interaction with the refractory period. Surrounding regions did not allow such a trade-off and could only trigger a single spike, after which the axon could not reset to fire again. In contrast, the unmyelinated axon’s spiking activity was maximized for an optimal sub-region of intermediate duty cycles where the firing rate approached the stimulus rate (PRF = 200 Hz). Interestingly, larger pressures offered a wider span of this optimal DC interval. Higher duty cycles (up to 100%) also generated spiking activity but also significantly interfered with the axon’s ongoing membrane dynamics and were thus less effective.

## Discussion

In this study, we used a novel computational framework to formulate several important predictions on the effects and mechanisms of US neuromodulation by intramembrane cavitation in peripheral fibers. First, single US pulses are capable of inducing *de novo* action potentials in both myelinated and unmyelinated peripheral axons. Second, these two fiber types share a common US recruitment mechanism: the stimulus onset induces a local drop in effective membrane capacitance at the acoustic focus and triggers passive depolarizing currents that raise charge density towards the spiking threshold. Third, while the two fiber types show a robust excitability across a wide range of carrier frequencies and acoustic pressure fields, they exhibit distinct sensitivities to temporal features of US stimuli. Myelinated axons exhibit a low (sub-millisecond) response latency due to their short membrane time constant and can therefore be excited by very short US pulses for which unmyelinated axons are unresponsive. However, for longer stimuli, unmyelinated axons can be excited at lower acoustic intensities than myelinated axons. Interestingly, this enhanced excitability in the rheobase regime is not caused by the absence of myelin, but rather, it is attributable to a smaller specific membrane capacitance 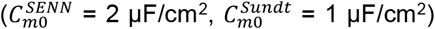. To the best of our knowledge, the biological origin of this capacitance difference remains undocumented. However, we found the magnitude ratio of this specific parameter to be conserved across a wide collection of biophysical models of myelinated and unmyelinated axons (Frankenhaeuser and Huxley, 1964; McIntyre et al., 2002; Tarnaud et al., 2018b), which supports the reliability of our conclusions. Fourth, the application of repeated US pulses induces a sustained spiking activity in both fiber types, the rate of which can be modulated by adjusting pulsing parameters. Particularly, myelinated axons robustly follow the stimulus rate over a wide range of PRFs, pulse durations and supra-threshold stimulus amplitudes, while unmyelinated axons show more complex dependencies on pulse durations / duty cycle and acoustic intensities. Moreover, both fiber types can be entrained into firing rates that are comparable to those resulting from electrical stimulation (Krauthamer and Crosheck, 2002), with myelinated axons showing a much higher upper limit (FR > 1 kHz) than their unmyelinated counterparts (FR < 180 Hz). The latter finding must be interpreted with caution, as the SENN myelinated axon model ignores subtle spiking adaptation phenomena and hence probably overestimates the physiological limit of the myelinated axon’s firing rate. Importantly, robust neuromodulatory effects can be obtained with both fiber types at relatively low duty cycles (DC < 50%) that prevent significant tissue heating. Together, these predictions define a comprehensive theoretical basis that can guide the design of US neuromodulation protocols.

### Applicability of the SONIC paradigm in multi-compartmental models

The SONIC paradigm relies on the assumption that membrane charge density and ion channel kinetics evolve at a much slower speed than microsecond-scale capacitance oscillations, thereby allowing for the accurate integration of neural responses using pre-computed cycle-averaged quantities of fast-oscillating variables. While that assumption is valid for point-neuron models (Lemaire et al., 2019), a recent study using a nanoscale two-compartment model have shown that under tight axial coupling conditions, strong intracellular currents mediate a significant intra-cycle charge redistribution that influences local membrane dynamics in a way that is not captured by the SONIC paradigm, resulting in overestimated sub-threshold charge build-ups and underestimated excitation thresholds (Tarnaud et al., 2020). It was also demonstrated that this numerical inaccuracy could be resolved by taking into account a limited number of Fourier components from precomputed oscillatory variables (as opposed to the SONIC approach that only considers their first component). Those findings raise legitimate concerns about the applicability of the SONIC paradigm in multi-compartmental models and prompted us to examine the conditions of its applicability, and whether it can be accurately used with the axon models of this study.

First, using a generic passive benchmark, we showed that SONIC accuracy is impacted by both intrinsic model properties and stimulus features, but also that this paradigm shows robust convergence if the underlying (membrane and axial) time constants of the considered neuron model are longer than the stimulus periodicity. Second, using axon-specific benchmarks, we demonstrated that the SONIC paradigm can accurately compute passive and active neural responses of both axon models of this study, across a vast majority of the LIFUS parameter space.

In the case of the unmyelinated axon, the axial time constant is a direct product of the spatial discretization of a continuous membrane 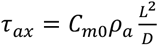, with a the compartment length). Hence, for small enough compartments, this time constant may become smaller than the stimulus periodicity and therefore sensitize the electrical system to intra-cycle variations. However, increasing the model resolution would also eliminate the spatial gradient in acoustic pressure across consecutive compartments, effectively eradicating the axial currents at the origin of SONIC divergence. In fact, the selected compartment length in this study (see Methods section) is in the order of 10-2 mm, i.e. already two orders of magnitude smaller than millimeter-scale pressure field variations.

Whether SONIC convergence can extend to other morphological models remains an open question, in particular as neurons of the CNS have a much slower membrane dynamics than peripheral axons (membrane time constants in the order of tens of milliseconds (Pospischil et al., 2008)) but possess tightly connected and heterogeneous morphological sections that may induce significant axial charge redistribution. In this case, a more computationally taxing approach considering extended Fourier decomposition might be required to achieve an acceptable level of accuracy.

### Generalizability of the morphoSONIC framework

Due to its intrinsic mechanoelectrical coupling, the NICE model is inherently tedious to simulate. In fact, capacitance oscillations induced by the mechanical membrane resonance introduce a high frequency capacitive displacement current 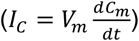 that greatly increases the associated numerical stiffness. We first observed that this stiffness could be reduced by recasting the differential system in terms of charge density (Lemaire et al., 2019). This strategy has since been employed in another study implementing the NICE model (Tarnaud et al., 2020), and is also at the core of the SONIC paradigm. Unfortunately, neither time-varying capacitance nor charge casting are natively supported by standard neuronal simulation environments such as NEURON. Consequently, computational studies on intramembrane cavitation have been implemented in custom software (Matlab or Python) and restricted to single and two-compartment models, partly because sub-optimal integration routines yield exorbitant simulation times and / or numerical instabilities for larger models.

Here, we derived a hybrid (charge and voltage casted) variant of the cable equation that is numerically reconcilable with both the NICE and SONIC paradigms and implemented it as an independent module that can be seamlessly integrated within the NEURON simulation environment. As such, the proposed approach provides a general solution to the problem of time-varying capacitance that is applicable to a wide variety of model types (single and double cable) and morphological structures (compartment number, branching patterns) seen across the central and peripheral nervous systems. Notably, this approach could also be used with enriched membrane mechanisms including lookup tables for additional Fourier components, as in (Tarnaud et al., 2020). Moreover, the choice of a NEURON-based implementation offers several advantages. First, it leverages NEURON’s optimized numerical integration pipelines while offering an appreciable abstraction level to the underlying differential systems. Second, it is applicable to a wide collection of biophysical models – as well as other resources – made available by the NEURON community (McDougal et al., 2017) with limited adaptation effort. Finally, although it has been used here with Gaussian field distributions approximating analytical solutions to simple physical problems, the morphoSONIC framework can easily be combined with finite-element-method (FEM) approaches. This refined multi-scale approach would enable the coupled simulation of complex acoustic propagation, pressure field distribution, and resulting neuronal responses inside anatomically accurate inhomogeneous tissue (such as the brain or the nerve environment).

### Comparison with empirical findings

As stated before, one of the major findings of this modeling study is that short US pulses are capable of inducing *de novo* action potentials in both myelinated and unmyelinated peripheral axons. This modeling prediction is in agreement with experimental observations from two recent studies showing that *in vivo* sonication of the mouse intact sciatic nerve directly activates myelinated fibers to induce motor responses (Downs et al., 2018), and that *ex vivo* sonication of unmyelinated crab leg nerve bundles generates compound action potentials (Wright et al., 2017). Interestingly, these studies reported significantly higher excitation thresholds (3.2 MPa and 1.8 MPa peak pressures around the fiber location for myelinated and unmyelinated axons, respectively) than the ones predicted here. Such differences could potentially arise from the intrinsic embedding of fibers within the neural tissue, increasing viscoelastic stresses on the membrane, and therefore hindering its mechanical resonance to acoustic perturbations (Krasovitski et al., 2011), a phenomenon that was not considered here. In fact, active neural responses in the extracted crab leg nerve bundles coincided with the presence of inertial cavitation in the surrounding medium, which may indicate higher thresholds for intramembrane cavitation in this specific environment. Nonetheless, considering that both studies employed minimal stimulus durations that fall within the fibers predicted rheobase regimes (4 ms and 8 ms for the myelinated and unmyelinated cases, respectively), the lower relative range of reported excitation thresholds for unmyelinated fibers corroborates our modeling predictions. Moreover, shorter response latencies were observed in myelinated fibers (Δt < 1 ms) than in unmyelinated fibers 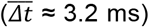, which is also in agreement with our findings. Significant variability in success rate and response latency was observed in the *ex vivo* crab leg nerve preparation, which is a departure from the deterministic nature of single fiber responses predicted by our current model. Nevertheless, the similarities in qualitative behavior between our theoretical results and these empirical observations provide a first indication that intramembrane cavitation could be a physiologically relevant US neuromodulation mechanism also in the peripheral nervous system. A more definitive answer to that question will require further experimental investigations, including a thorough comparison of excitation thresholds across fiber types and diameters within the same nerve environment and across a wide range of pulse durations and acoustic beam widths.

### Therapeutic implications

Beyond mechanistic investigation, our findings further emphasize the potential of LIFUS as a noninvasive neuromodulation technology and its applicability to peripheral structures. In fact, we predict that LIFUS can be used to robustly modulate the spiking activity of both myelinated and unmyelinated fibers, meaning that it could be used to encode sensory information or elicit motor responses. In this context, the lack of clear dependency of LIFUS excitation thresholds on fiber diameter represents a disadvantage, as it excludes the possibility to discriminate across different populations of myelinated fibers and, hence, to target a specific peripheral pathway. However, the ability to selectively target unmyelinated C-fibers, which carry pain and temperature afferent signals, ushers in the possibility to encode new types of sensory information in artificial limbs without interfering with other haptic, i.e. tactile (Petrini et al., 2019; Raspopovic et al., 2014; Valle et al., 2018) and proprioceptive (D’Anna et al., 2019), modalities. To the best of our knowledge, this feature has never been achieved with standard electrical stimulation techniques. The encoding of temperature information would be particularly interesting to enrich the sensory feedback in neuroprosthetic devices and improve user experience (Mendez et al., 2020).

## Conclusions

In this study, we present a novel computational framework to investigate the mechanisms of US neuromodulation by intramembrane cavitation in morphologically realistic neuron models, using the NEURON simulation environment. The new framework is used to predict cell-type-specific responses of myelinated and unmyelinated peripheral axons to acoustic pressure fields. These predictions are in qualitative agreement with recent empirical observations, and open new avenues for the use of US as a neuromodulation technology in the peripheral nervous system. Yet, closer quantitative comparison with experimental data will be necessary to further validate or reject the underlying mechanism. Furthermore, we plan to couple our modular framework with acoustic propagation models to formulate more detailed predictions of neural responses upon sonication by realistic acoustic sources and to inform the development of application-specific ultrasonic devices.

## Materials and Methods

### The NICE model

The NICE electromechanical model developed by Plaksin et al. (Plaksin et al., 2014) provides a mathematical formulation of the intramembrane cavitation hypothesis. Mechanically, the periodic cavitation of a single bilayer sonophore is described by two differential variables: the deflection of a leaflet apex from its resting position in the transmembrane plane (*Z*) and the internal gas content in the sonophore cavity (*n_g_*). The resting leaflet position results from a pressure balance between several static pressure forces, namely the elastic tension developing in the leaflets (*P_S_*), attractive and repulsive intermolecular forces between leaflets (*P_M_*), internal gas pressure in the sonophore cavity (*P_G_*), the electrical pressure resulting from the membrane polarity (*P_Q_*), and a constant hydrostatic term (*P_0_*). Upon perturbation by a time-varying acoustic pressure *P_A_(t)*, the dynamic pressure imbalance drives a normal acceleration that deforms the leaflets in antiphase, generates viscous forces in the membrane (*P_VS_*) and surrounding medium (*P_VL_*), and triggers gas transport across the cavity. These oscillatory dynamics are captured by the following differential system (all pressure terms and parameters are defined in (Lemaire et al., 2019)):

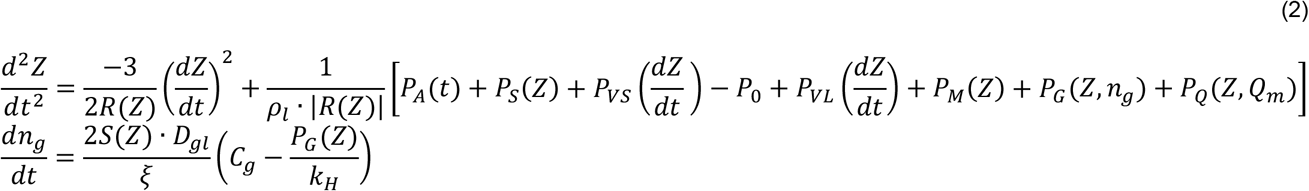

Electrically, the development of an electrical response across the membrane is captured by a modified Hodgkin-Huxley differential system, describing the evolution of the membrane charge density (*Q_m_*) as the negative sum of voltage-dependent ionic currents with specific conductances *gi* and reversal potentials *Ei*. In this system, time-varying ionic conductances are the product of one or multiple gating variables (*x,* with *x* ∈ {*m*, *h*, *n*, *p*, …}), whose evolution is regulated either by a voltage-dependent activation and inactivation rate constants (*αx* and *βx*, respectively) or by a steady state probability *x_∞_* and a time constant *τ_x_* (also both voltage-dependent), yielding the following system (note that charge-casting was introduced in (Lemaire et al., 2019)):

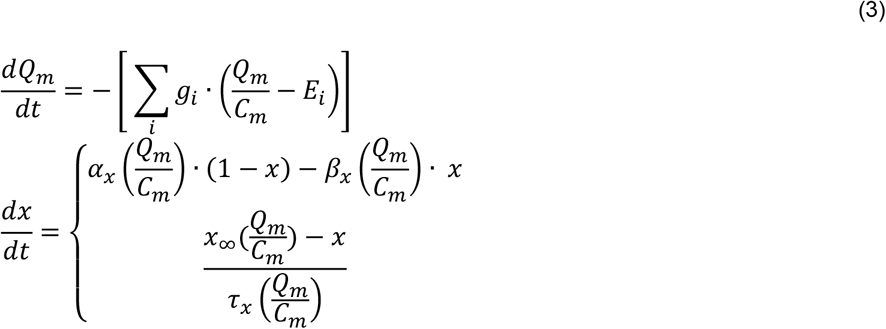

The coupling between these two systems is modelled by a bi-directional piezoelectric effect. Mechano-electrical transduction results from the periodic deflections of the sonophore leaflets, inducing high frequency oscillations in the local membrane capacitance (given by 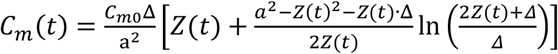, as in (Plaksin et al., 2014)).

Considering a larger, macroscale portion of membrane area, local fluctuations of membrane capacitance around individual sonophores influence the spatial average of membrane capacitance, calculated as a weighted mean of the resting and dynamic capacitances (*C_m_* = *C_m_*(*t*)*f_s_* + *C_m0_*(1 − *f_s_*), where *f_s_* is the sonophore membrane coverage fraction). This global fluctuation then causes large-amplitude oscillations of the transmembrane potential in the compartment of interest (*V_m_* = *Q_m_*/*C_m_* in equation (3)). Reversibly, electro-mechanical transduction results from progressive changes in the membrane electrical polarity that dynamically modify the electrical pressure exerted on the sonophore leaflets and the resulting pressure balance (*P_Q_* in equation (2)), thereby influencing the sonophore cavitation dynamics.

### The SONIC model

The SONIC model (Lemaire et al., 2019) uses temporal multiscaling to separate the two relevant time scales of the NICE model, namely microsecond-scale mechanical oscillations and millisecond-scale development of neuronal responses. It is based on the observation that ion channel gates – whose time constants are typically in the millisecond range – do not follow large-amplitude, high-frequency variations of transmembrane potential observed in the NICE model, but rather adapt to the temporal average of voltage oscillations over an acoustic cycle. As a result, the evolution of membrane charge density and ion channels gating variables can be expressed as a function of an effective membrane potential (*V_m_\**) and effective activation and inactivation rate constants (*α \** and *β \**, respectively, for each gating variable *x*), representing the average value of their original, voltage-dependent counterparts (*V_m_*, *α_x_* and *β_x_*, respectively) over an acoustic cycle:

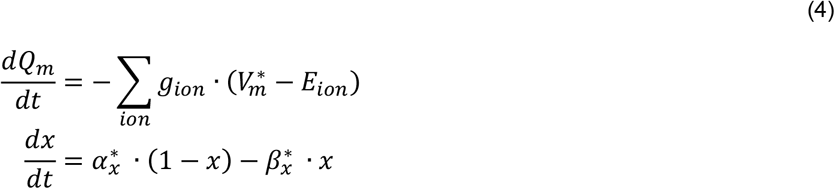

The SONIC model uses a sequential approach to compute electrical responses of a given neuron type to various LIFUS stimuli. First, a parallelized precomputation step is performed (once per neuron type) in which the mechanical system (equation (2)) is simulated for various combinations of sonophore radii (*a*), US frequencies (*fUS*), acoustic peak pressure amplitudes (*A_US_*), and membrane charge densities, covering the LIFUS parametric space, sonophore geometrical range and membrane physiological range. Each simulation is run until a limit cycle is detected, at which point, the profile of oscillating membrane capacitance over the last acoustic cycle is extracted, dampened for a range of sonophore membrane coverage fractions, and used to compute effective variables stored in multi-dimensional lookup tables. Second, the electrical response of the neuron to a given LIFUS stimulus is rapidly computed at runtime by interpolating effective variables at (*a, f_s_, f_US_, A_US_*) and (*a, f_s_, f_US_, 0*) to yield 1D projected vectors in the *Q_m_* space, which are then used to interpolate effective variables and solve equation (4) during US-ON and US-OFF periods, respectively.

### A hybrid multi-compartmental, multi-layer electrical circuit

In its most basic form, the multi-compartmental expansion of point-neuron NICE / SONIC models requires the addition of axial current terms contributing to the evolution of charge density in each compartment (see (Lemaire et al., 2019), equation 5). However, that formulation only considers intracellular axial coupling, and is therefore not adapted to “double-cable” models that account for both intra and extracellular longitudinal coupling. More importantly, the use of explicit current terms representing axial coupling is prone to yielding numerical instabilities in the presence of tightly connected sections or abrupt changes in voltage gradients. Hence, in this study, we derived a hybrid multi-compartmental multi-layer electrical circuit that is applicable to both myelinated and unmyelinated structures with temporally and spatially varying membrane capacitances, and compatible with reference numerical integration schemes and simulation environments.

The circuit model is composed of multiple longitudinal compartments, each represented by a pair of intracellular and extracellular voltage nodes (*Vi* and *V_x_*, respectively) on either side of the plasma membrane with time-varying capacitance *C_m_(t)*. The voltage difference across the plasma membrane *V_m_\** = *V_i_* - *V_x_* influences the opening and closing of distinct ion channels, ultimately giving rise to a net membrane ionic current *I_ion_*. On the extracellular side, a transverse resistor-capacitor (RC) circuit of conductance *g_x_* and capacitance *C_x_* represents the myelin membrane and connects the extracellular node to the extracellular driving voltage *E_x_*, which is usually grounded but can also have a value imposed by an external electrical field. Longitudinally, neighboring nodes are connected intracellularly and extracellularly by axial conductors (*G_a_* and *G_p_*, respectively). All variables and parameters of the circuit are described in **Table 1**.

**Table 1.** Parameters and variables of the hybrid multi-compartmental, multi-layer electrical circuit.

| Parameter / Variable | Symbol | Unit |
| --- | --- | --- |
| Intracellular voltage | $V_i$ | mV |
| Extracellular voltage | $V_x$ | mV |
| Transmembrane voltage | $V_m$ | mV |
| Transmembrane charge density | $Q_m$ | nC/cm<sup>2</sup> |
| Extracellular driving voltage | $\Phi_e$ | mV |
| Membrane capacitance | $C_m$ | μF/cm<sup>2</sup> |
| Intracellular stimulation current | $I_s$ | mA/cm<sup>2</sup> |
| Net transmembrane ionic current | $I_{ion}$ | mA/cm<sup>2</sup> |
| Intracellular axial conductance | $G_a$ | S |
| Capacitance of surrounding extracellular membrane (e.g. myelin) | $C_x$ | μF/cm<sup>2</sup> |
| Transverse conductance of surrounding extracellular membrane (e.g. myelin) | $g_x$ | S/cm<sup>2</sup> |
| Extracellular axial conductance (e.g. periaxonal space) | $G_p$ | S |
| Membrane area of the compartment | $A_m$ | cm<sup>2</sup> |

For any compartment *k* connected to a set of neighboring compartments, the application of Kirchhoff’s law at the corresponding intracellular and extracellular nodes yields the following current balance equations:

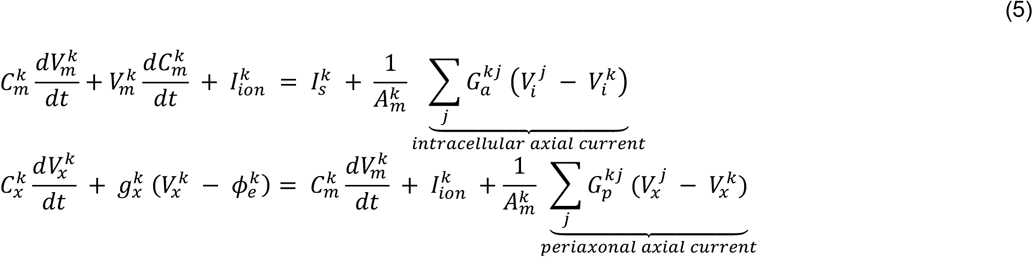

Using *V_i_* = *V_m_* + *V_x_*, and re-arranging the terms, we find:

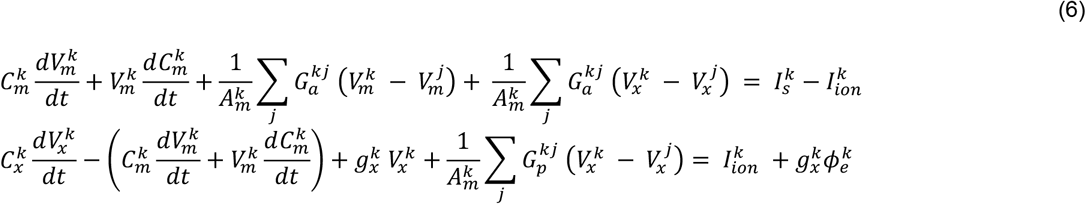

By substituting transmembrane voltage for transmembrane charge density 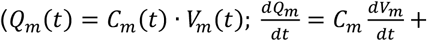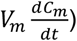 and defining 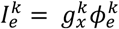 as the extracellular driving current, we obtain:

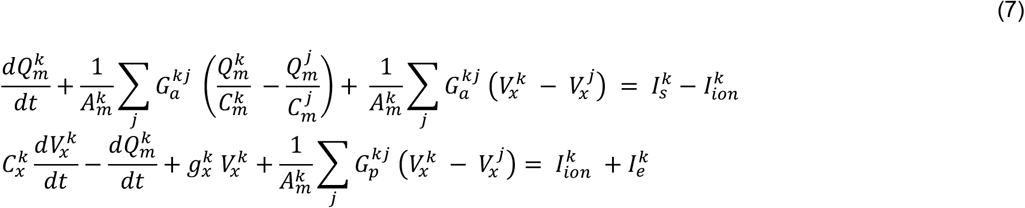

By applying the above equations to a model of *n* compartments connected in series, we obtain a hybrid charge-voltage partial differential equation system of size *2n* that can be described as:

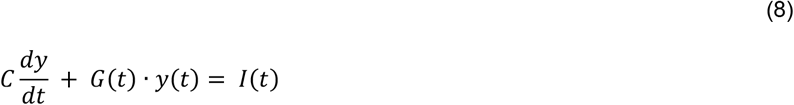

where:

- *y* is a hybrid vector of transmembrane charge density and extracellular voltage, and *dy/dt* its temporal derivative;
- *C* is a constant matrix composed of both capacitance terms (multiplying voltage elements of *dy/dt*) and “identity” terms (multiplying charge elements of *dy/dt*);
- *G(t)* is a time-varying matrix composed of both conductance terms (multiplying voltage elements of *y*) and “frequency” terms (conductance by capacitance ratios in MHz, multiplying charge elements of *y*); and
- *I(t)* is a time-varying vector of stimulation and membrane currents

This matrix formulation allows for the use of implicit methods to solve the differential equation problem, thus providing an enhanced stability over explicit schemes.

Moreover, by mapping the first *n* elements of the *y* vector to transmembrane charge density nodes and the following *n* elements to extracellular voltage nodes, we can describe the *C*, *G* and *I* terms of the system as combinations of block matrices and vectors, i.e.:

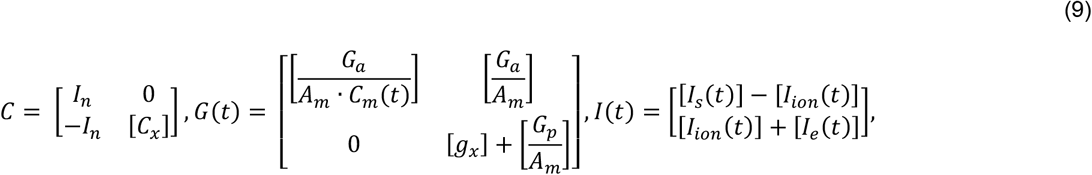

where:

- *I_n_* is an n-by-n identity matrix;
- [*C_x_*] is an n-by-n diagonal matrix of transverse extracellular membrane (e.g. myelin) capacitance;
- 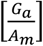 and 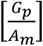 are n-by-n tridiagonal matrices of intracellular and extracellular axial conductance, respectively, where each row is normalized by the corresponding node’s membrane area;
- 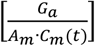 is an n-by-n tridiagonal matrix of intracellular axial conductance where each row is normalized by corresponding node’s membrane area and each column is dynamically normalized by the time-varying membrane capacitance of the corresponding node;
- [*g_x_*] is an n-by-n diagonal matrix of transverse extracellular membrane (e.g. myelin) conductance; and
- [*I_s_* (*t*)], [*I_ion_* (*t*)] and [*I_e_* (*t*)] are n-sized, time-varying vectors of intracellular stimulation currents, transmembrane ionic currents and extracellular driving currents, respectively.

We implemented this hybrid system in NEURON (Hines and Carnevale, 1997), a reference computational environment for neuronal simulations that uses a very similar matrix formulation to enable numerical integration by implicit schemes. However, since that environment is not designed for models of varying capacitance or for hybrid charge-voltage casting, we employed three main adaptation strategies. First, a unit capacitance was set to all membrane mechanisms, thereby implicitly setting the *I_n_* upper block matrix and effectively transforming NEURON’s internal variable *v* as an alias to transmembrane charge density. Second, pressure amplitude and charge density dependent lookup tables of effective SONIC terms (transmembrane potential and ion channels rate constants obtained from original SONIC lookup tables (Lemaire et al., 2019)) were dynamically inserted into these mechanisms to compute the evolution of voltage, ion channels states and ionic currents via bilinear interpolation (thereby implicitly setting the *I_ion_* upper block vector). Third, a “Linear Mechanism” object was implemented that defines alternative *C’*, *G’* and *I’* terms to complete the hybrid circuit setup once the model’s compartments and their connections are defined:

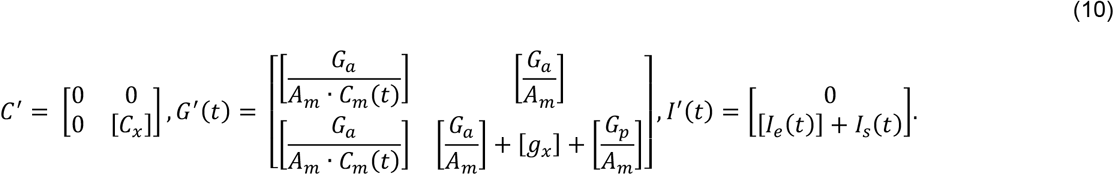

It should be noted that the terms −*I_n_* and [*I_ion_*(*t*)] in the lower block are replaced by equivalent axial conduction and intracellular stimulation current terms (using the equality of the upper block) to remove the need to access the net membrane current (a hidden NEURON variable). Numerical integration is then carried out by NEURON’s embedded general sparse matrix solver (a differential-algebraic solver with a preconditioned Krylov method from the SUNDIALS package (Hindmarsh et al., 2005)) using a variable time step with a pure absolute error tolerance criterion (∈ = 10^−3^), while dynamically updating *C_m_*-dependent terms in the *G’* matrix throughout the simulation. Compared to previous approaches using explicit axial current terms (Lemaire et al., 2019), this implicit integration scheme offers increased numerical stability.

### Myelinated and unmyelinated morphological axon models

We used single-cable axonal representations allowing for a numerically valid incorporation of the SONIC paradigm while maintaining a certain level of morphological realism.

Our myelinated axon model was based on the spatially-extended nonlinear node (SENN) model developed by Reilly et al. (Reilly et al., 1985). This model represents myelinated axons as a set of nodes with active membrane dynamics based on the Frankenhaeuser-Huxley equations for a Xenopus Ranvier node (Frankenhaeuser and Huxley, 1964) including fast sodium (I_Na_), delayed-rectifier potassium (I_Kd_), non-specific delayed (I_P_) and non-specific leakage (ILeak) currents, connected by intracellular resistors representing the myelinated internodes (**Figure 1A-B**). This representation omits specific features of myelinated axons (namely transmembrane internodal dynamics and extracellular longitudinal coupling), but it incorporates enough morphological complexity to provide quantitatively accurate predictions of their excitability by electrical fields. In fact, it represents a standardized model for electromagnetic exposure safety assessment (Reilly, 2011). Our unmyelinated axon model was based on the work of Sundt et al. (Sundt et al., 2015), representing the continuous unmyelinated neurite as a set of nodes containing fast Sodium (I_Na_), delayed-rectifier Potassium (I_Kd_), and leakage (I_Leak_) membrane currents, also connected by intracellular resistors (**Figure 1C-D**).

The selected axon models were validated numerically by verifying specific physiological features (spike amplitude, conduction velocity, threshold excitation current for various pulse widths) against the reference literature (Reilly et al., 1985; Sundt et al., 2015), using NEURON’s native voltage-based connection scheme with constant membrane capacitance. For the unmyelinated model, a convergence study was carried out to determine the optimal spatial discretization. Unmyelinated compartments were progressively and uniformly shortened from 1 mm to 5 μm, and an optimal segment length was defined as the maximal length for which all physiological features were within 5% of their converging values (obtained for the shortest segment length). As the optimum segment length exhibited a clear dependency on fiber diameter, we performed a piecewise linear fit within the 0.5 – 1.5 μm range to obtain a fiber diameter-dependent formulation: *Lopt* = min(16.4 μm^−1^ · *D_fiber_* + 9.1 μm; 22 μm). Finally, we validated our hybrid circuit implementation by comparing direct voltage traces, as well as physiological features, to those obtained with the “native” implementation.

Membrane equations of both models were adapted to 36°C by applying a Q10 correction with a factor of 3 (as in (Sundt et al., 2015)), and lookup tables of SONIC effective variables were generated for the membrane circuits of both models to enable their simulation upon acoustic perturbations (**Figure 1E**).

Beyond the differences in morphologies and ion channel populations, the two models have different resting membrane capacitances 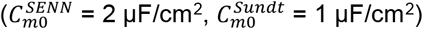, thereby implying variations in their charge density by a factor of 2 for identical voltages. Therefore, to allow for an unbiased comparison of electrical responses across the two models, charge-density based criteria and derived outputs were divided by the resting capacitance of the associated model, referred to as the “normalized charge density” (in mV).

### Modeling of exogenous acoustic and electrical stimuli

In order to evaluate the effect of exogenous electrical and ultrasonic stimulation on isolated fibers, we modeled the propagation of both electrical and acoustic fields from a realistic remote excitation source to the target through a homogenous intraneural medium. To this end, we considered a 3-dimensional (*x*, *y*, *z*) coordinate system in which the fiber was aligned on the *x* axis and centered at the origin.

For ultrasonic stimulation, we considered a single-element planar acoustic transducer with a center in the *yz* plane and a normal vector along the *z*-axis, and a homogenous, water-like propagation medium (density *ρ* = 1000 kg/m3, speed of sound *c* = 1500 m/s). We modeled acoustic distribution in the *yz* propagation plane using the Distributed Point Source Method (DPSM) (Yanagita et al., 2009), which provides accurate approximations of the Rayleigh-Sommerfeld integral (RSI) in homogenous medium. That is, assuming a uniform particle velocity normal to the transducer surface of amplitude *v*_0_, the acoustic peak pressure amplitude at each field point (*x*, *z*) for an acoustic frequency *f* can be computed as:

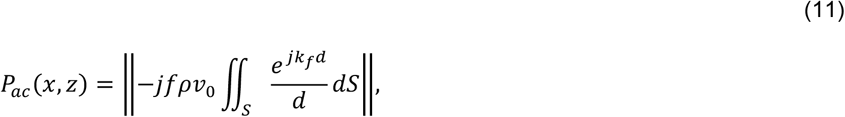

where *j* is the unit imaginary number, *k_f_* = 2*πf*/*c* is the wave number, and 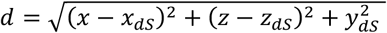 is the distance between the field point and a surface element *dS*. We numerically approximated this integral as the sum of individual contributions of a finite set of *M* uniformly distributed point sources – each associated with a surface area Δ*S* arranged in a concentric fashion on the transducer surface:

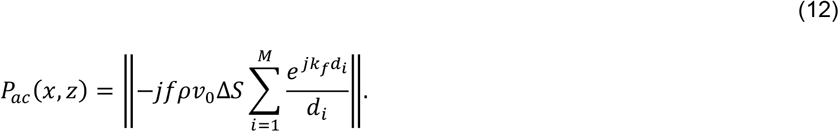

Here again, we performed a sensitivity analysis to determine the optimal density of point sources required to achieve a good prediction accuracy. Starting with a low source density (10 samples / mm^2^), the predicted pressure distribution along the central *z* axis was evaluated against the corresponding closed form RSI solution 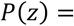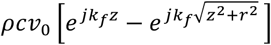, with *r* the transducer radius), and source density was increased until the variation of the root mean square error (RMSE) fell below a threshold value (10 kPa). We then selected the minimal value satisfying that criterion over a wide frequency range (500 kHz – 5 MHz), yielding an optimal density of 217 samples / mm^2^.

Finally, we evaluated pressure distributions along the transverse *x* axis at the acoustic focal distance (calculated as 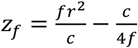 for each combination of transducer radius and US frequency.

For electrical stimulation, we considered a point source electrode located in the *xz* plane and an anisotropic conductivity tensor characteristic of the mammalian endoneurium (longitudinal resistivity *ρ*x = 175 Ω ∙ *cm*, transverse resistivity *ρ_yz_* = 1211 Ω ∙ *cm*) (Ranck and Bement, 1965). Extracellular potentials at each field point (*x*, *z*) were computed with the formula:

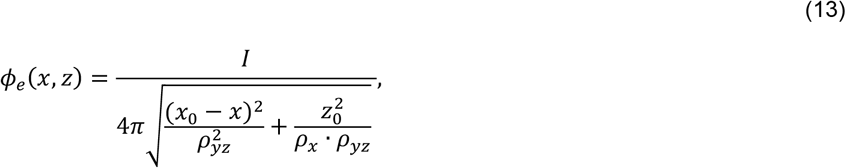

where *I* is the injected current and (*x_0_*, *z_0_*) are the electrode coordinates, and equivalent sets of intracellular currents were used to simulate the influence of the extracellular electric field, as in (McIntyre et al., 2002).

Note that equations (12) and (13) provide closed-form expressions to predict the qualitative nature of ultrasonic and electric field distributions along a fiber, thereby allowing general trends about the impact of those distributions on axon excitability to be established. However, they only consider propagation within a homogeneous medium, which is a limitation.

### Two-compartment SONIC benchmarks

The passive benchmark model was composed of two passive compartments with identical geometries and membrane properties (Cm0 = 1 μF/cm^2^, V_m0_ = E_Leak_ = −70 mV) connected by an intracellular conductance G_a_. Membrane and axial conductances were mapped to equivalent time constants (τ_*m*_ = *C_m0_*/*g_Leak_* and τ_*ax*_ = *C_m0_A_m_*/*G_a_*, respectively) allowing a direct comparison with the stimulus periodicity. Simulation durations were fixed to five times the longest time constant for each combination in order to ensure steady-state convergence, while ensuring at least 10 acoustic cycles.

The axon benchmark models were composed of two identical compartments with axon-specific morphological properties and full membrane dynamics of each fiber type. Physiologically relevant simulation durations known to elicit spiking activity in each model (1 ms and 10 ms for the myelinated and unmyelinated cases, respectively) were used.

All benchmark models were simulated under the NICE and SONIC paradigms using frequency-dependent time steps in both cases (dt_SONIC_ = 1 / f_US_, dt_NICE_ = 0.01 / f_US_). For each condition, the accuracy of the SONIC paradigm was evaluated by computing the maximal root mean square error (RMSE) between normalized charge density profiles of the SONIC solution and the cycle-averaged NICE solution (*∈_max_*, in mV).

## Acknowledgments

This work was partly funded by the Wyss Center for Bio and Neuroengineering (https://www.wysscenter.ch/) and by the Bertarelli Foundation (https://www.fondation-bertarelli.org/). The funders had no role in study design, data collection and analysis, decision to publish, or preparation of the manuscript.

## Author contributions

T.L. conceptualized the study, implemented the models, performed simulations, analyzed the results, prepared the figures and wrote the manuscript. E.V. performed simulations, analyzed results and reviewed the manuscript. E.N. conceptualized the study, analyzed results and reviewed the manuscript. N.K. reviewed the manuscript. S.M. supervised the study and reviewed the manuscript.

---

**Figure S1.**
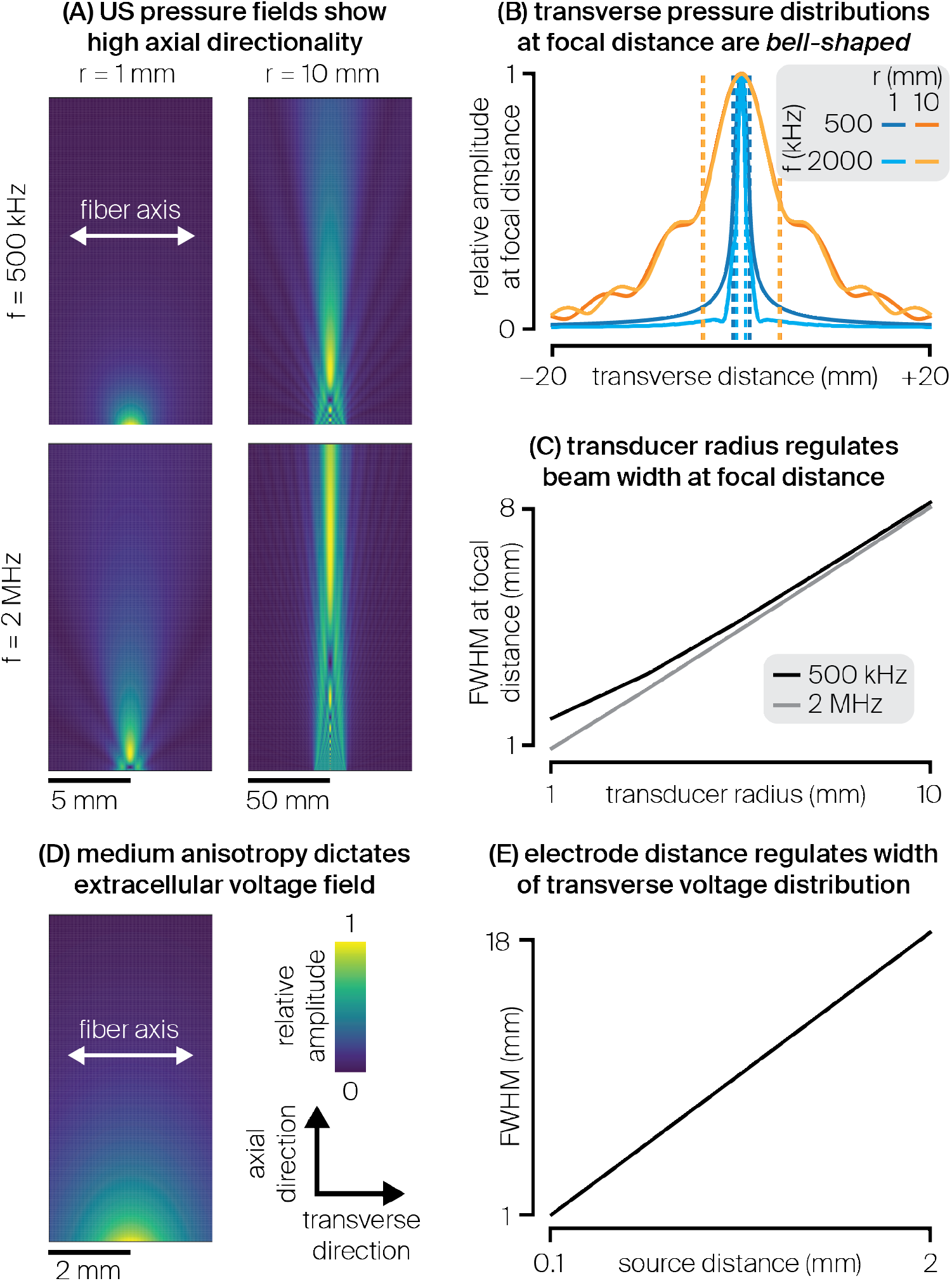
Qualitative nature of exogenous acoustic and electrical fields. **(A)** Normalized two-dimensional acoustic pressure amplitude distribution across the propagation plane computed upon sonication by a single-element planar transducer immersed in water-like medium, for various combinations of transducer radius and US frequency. A white line indicates the fiber’s axis as considered in this work. **(B)** Normalized transverse pressure distribution measured at the transducer’s focal distance, for the same combinations of transducer radius and US frequency. Dotted lines indicate FWHMs for each distribution. **(C)** FWHM of the pressure amplitude distribution along the fiber axis as a function of the transducer radius, for two characteristic US frequencies. **(D)** Normalized two-dimensional voltage distribution across a two-dimensional plane generated by a point-source electrode placed in a nerve-like anisotropic medium. A white line indicates the fiber’s axis as considered in this work. **(E)** FWHM of the extracellular voltage distribution along the fiber axis as a function of the electrode-fiber distance.

